# Targeting the YAP1/TAZ-TEAD Transcriptional Axis Reveals a Therapeutic Vulnerability in Androgen Receptor Signaling Inhibitor (ARSI)-Resistant Prostate Cancer

**DOI:** 10.64898/2026.11.19.767854

**Authors:** Vandana Mohan, Santosh Lamichhane, Praveen Kumar Jaiswal, Guobin Kang, Dorota D. Wyczechowska, Sweaty Koul, Hari K Koul

**Author notes:** Corresponding Author: Hari K. Koul, MSc., PhD, Department of Interdisciplinary Oncology, Louisiana Cancer Research Center Louisiana State University Health Sciences Center, New Orleans, LA 70112, USA Email: [ ].

## Abstract

Therapeutic resistance remains a major barrier in advanced prostate cancer (PCa), particularly after prolonged treatment with second-generation androgen receptor signaling inhibitors (ARSIs) such as enzalutamide. Tumors can evade these drugs by escaping androgen receptor (AR) dependence through cellular plasticity and lineage transitions, but alternative resistance networks remain poorly understood. Enzalutamide-resistant (ENZ-R) PCa cells showed marked activation and nuclear enrichment of the YAP-1/TAZ-TEAD complex. Disrupting this transcriptional interface with CA3 shows strong anti-metastatic effects, inhibiting migration, invasion, and colony formation across AR-dependent and AR-independent PCa cell lines, suggesting efficacy regardless of cellular lineage or AR status. Similar effects with GNE-7883, K-975, and siRNA knockdown supported a class effect. In vivo, CA3 treatment substantially reduces tumor burden in PC3 and DU145 xenograft models, supporting the systemic efficacy of disrupting the YAP-1/TAZ-TEAD axis in aggressive, AR-independent disease. Mechanistically, this anti-tumor response is driven by potent cytotoxicity: CA3 induces substantial DNA damage and apoptosis in both in vitro and in vivo settings. Moreover, we identified increased YAP-1/WWTR1(Taz) expression as a negative predictor of therapeutic response in PCa patients. These findings indicate that enzalutamide resistance creates a targetable dependence on functional Hippo transcription. Small-molecule disruption of YAP-1/TAZ-TEAD may offer a rational strategy to counter treatment resistance and lineage escape in advanced PCa.

**YAP-1/TAZ-TEAD inhibition via CA3 reverses Enzalutamide resistance and suppresses tumorigenesis in advanced prostate cancer**

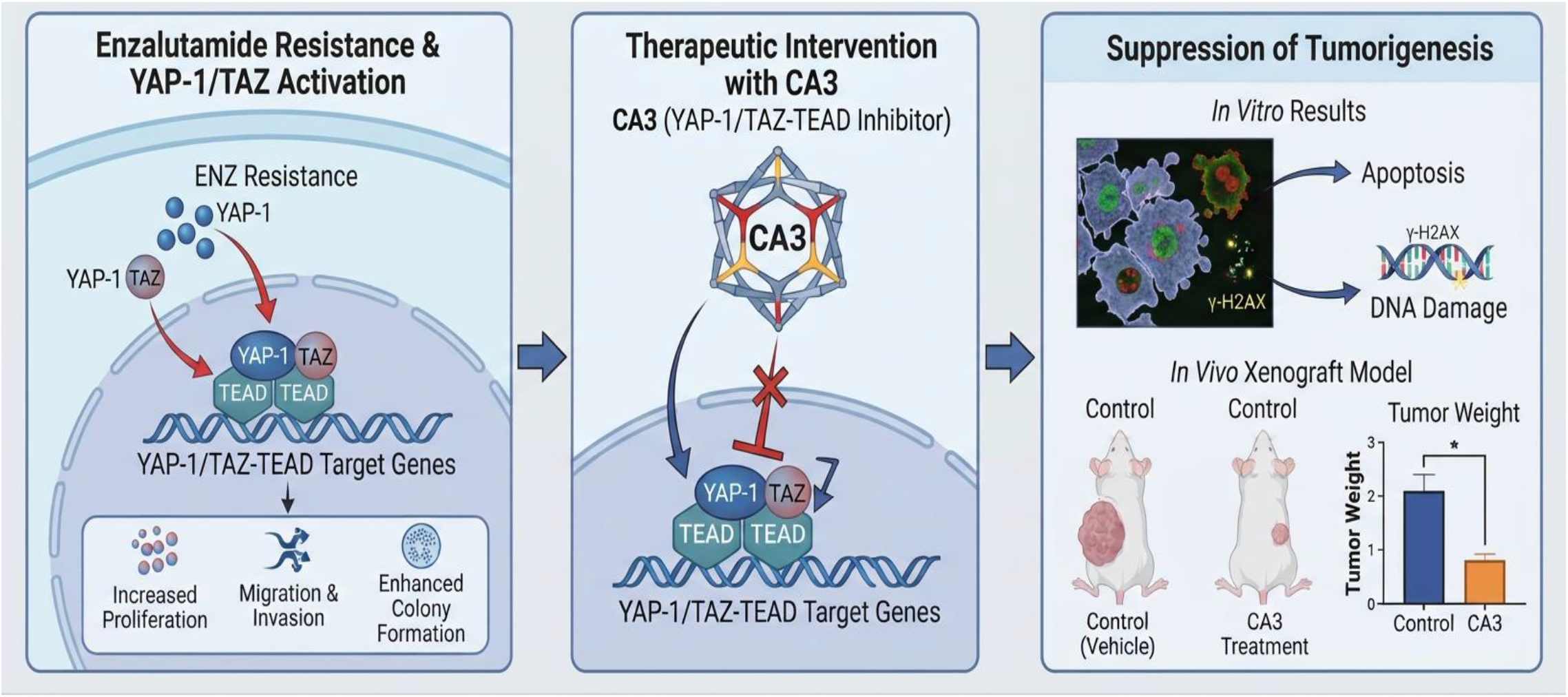

Microenvironmental stress and therapeutic pressure enrich nuclear YAP-1/TAZ-TEAD transcriptional activity, accelerating cell proliferation, migration, invasion, and robust colony formation in enzalutamide-resistant prostate cancer cells. Targeted intervention with the small-molecule inhibitor CA3 effectively blocks this signaling axis, disrupts nuclear complex interactions, and significantly reduces tumor weight and overall burden in aggressive in vivo xenograft models. Moreover, CA3 treatment induces robust apoptosis and DNA damage, as evidenced by increased γ-H2AX expression, highlighting the Hippo-YAP-1/TAZ pathway as a critical therapeutic vulnerability for countering advanced anti-androgen resistance. (*Graphical abstract image was generated using Google AI*)

**Highlights:**

1. Hippo-YAP/TAZ-TEAD activity is strongly enriched in enzalutamide-resistant PCa
2. CA3 disrupts YAP-1/TAZ to suppress PCa cell growth, motility, and clonogenicity.
3. Systemic CA3 treatment reduces tumor burden and drives microenvironment remodeling.
4. WWTR1 (TAZ)/YAP-1 expression is a prognostic marker of poor survival in prostate cancer patients.
5. Targeting YAP-1/TAZ-TEAD triggers DNA double-strand breaks and impairs RAD51 repair, highlighting potential vulnerability in ASI-resistant PCa.

## Introduction

Prostate cancer (PCa) remains the second most commonly diagnosed malignancy and the fifth leading cause of cancer-related death among men worldwide ^1,2^ . Although localized, early-stage disease responds well to initial treatment, advanced PCa can develop an aggressive, systemic phenotype ^3^ . Androgen deprivation therapy (ADT) is the established first-line treatment for advanced disease, but tumors almost universally evade this approach over time, progressing to lethal castration-resistant prostate cancer (CRPC) ^4,5^ . The introduction of highly potent, second-generation androgen receptor signaling inhibitors (ARSIs), particularly enzalutamide (ENZ) and abiraterone acetate, transformed CRPC management by greatly extending overall and progression-free survival^6,7^.

Despite these clinical milestones, prolonged, intensive suppression of the AR axis creates evolutionary selective pressure that drives diverse mechanisms of therapeutic resistance. Classic resistance mechanisms often involve AR-centric alterations, including AR amplification, gain-of-function mutations, and expression of constitutively active splice variants such as AR-V7. These alterations sustain oncogenic signaling despite maximal ARSI blockade ^8,9^ . However, as clinical strategies suppress AR signaling more profoundly, tumors increasingly bypass AR dependence altogether by exploiting non-AR-driven survival networks ^10^ .

A key mechanism underlying this AR-independent evasion is cellular plasticity and dedifferentiation ^11^ . In response to modern ARSIs, prostate adenocarcinoma cells can lose their committed, androgen-dependent luminal epithelial identity and revert to a primitive, multipotent, stem-like state ^12^ . This dedifferentiation gives tumor cells remarkable phenotypic flexibility, enabling them to dynamically remodel their transcriptional and epigenetic landscapes ^13^ . One major clinical manifestation of this plasticity is the lineage transition from adenocarcinoma to neuroendocrine prostate cancer (NEPC) ^14^ . Although de novo neuroendocrine prostate tumors are exceedingly rare (<2% of primary presentations), therapy-induced lineage plasticity accounts for approximately 17–30% of advanced disease progressions following ARSI exposure ^15,16^ . Together, these fluid cellular states pose a major clinical hurdle, allowing tumors to respond to changing microenvironmental and therapeutic landscapes ^17^ .

Emerging evidence identifies dysregulation of the Hippo signaling pathway as a key driver of multiple therapeutic adaptations, cellular dedifferentiation, and lineage escape ^18,19^ . When the core upstream Hippo kinase cascade is inactivated or bypassed, the downstream transcriptional co-activators YAP-1 (YAP-11) and TAZ (WWTR1) can translocate freely into the nucleus ^20^ . There, they physically interact with TEAD family transcription factors (TEAD 1–4) to activate a robust oncogenic gene expression program that regulates cellular survival, stemness, epithelial-to-mesenchymal transition (EMT), and cellular plasticity ^21,22^ . Importantly, elevated nuclear YAP-1/TAZ-TEAD activity provides a functional bridge to an AR-independent state. It correlates not only with neuroendocrine differentiation markers but also with broader intrinsic and acquired resistance to next-generation anti-androgens ^23^ . Moreover, YAP-1/Taz expression has been shown to be enriched in a stem-cell-like phenotype in prostate cancer ^24^ . Consistent with these findings, our results show that YAP-1/TAZ-TEAD transcriptional activity is markedly enriched in enzalutamide-resistant PCa cells, and that high WWT1 expression is inversely associated with prostate cancer patient survival, supporting its role as an alternative survival pathway under chronic ARSI treatment.

Given the importance of this adaptive loop across multiple resistant phenotypes, pharmacologically targeting the YAP-1/TAZ-TEAD interaction has emerged as a promising therapeutic strategy ^25^. In this study, we assessed the therapeutic potential of the small-molecule YAP-1/TAZ-TEAD inhibitor CA3 in advanced PCa models. We show that inhibiting YAP-1/TAZ-TEAD efficiently suppresses proliferation in both parental and enzalutamide-resistant PCa cells. CA3 also shows strong anti-metastatic effects, inhibiting migration, invasion, and colony formation across AR-dependent and AR-independent PCa cell lines, suggesting efficacy regardless of cellular lineage or AR status. In vivo, CA3 treatment substantially reduces tumor burden in PC3 and DU145 xenograft models, supporting the systemic efficacy of disrupting the YAP-1/TAZ-TEAD axis in aggressive, AR-independent disease. Mechanistically, this anti-tumor response is driven by potent cytotoxicity: CA3 induces substantial DNA damage and apoptosis in both in vitro and in vivo settings. Moreover, we identified increased YAP-1/WWTR1(TAZ) expression as a negative predictor of therapeutic response in PCa patients. Taken together, our results and preliminary findings ^26,27^ show that targeting the YAP-1/TAZ-TEAD transcriptional complex with small molecules such as CA3 offers a rational therapeutic approach to counter ARSI resistance in Prostate cancer.

## Material and Methods

Reagents: All cell culture media were purchased from Corning. Trypsin with 1 mM EDTA (SM-2003-C) was purchased from Sigma-Aldrich. Antibodies for YAP-11 (12395), P-YAP-11 (13008), TAZ (#72804), and TEAD1 (12292) were purchased from Cell Signaling. TEAD4 (1241-8-1-AP) was purchased from Protein Tech (Thermo Fisher). The TEAD2 (PAS-116018) antibody was purchased from Invitrogen. HRP-conjugated anti-mouse IgG (115-035-003) and anti-rabbit IgG (111-035-003) were obtained from Jackson ImmunoResearch. Cell culture inserts (8.0 μm) (353097), Matrigel for in vitro assays (354234), and Matrigel (354230) for in vivo experiments were procured from BD Biosciences. Enzalutamide (HY70002) was purchased from MedChemExpress. The inhibitor CA3 (#S8661) was purchased from Target Mol, dissolved in DMSO to prepare a stock solution according to the inhibitor manual, and used at the desired concentration in the experiments.

### Cell culture

Human Prostate cancer cell lines PC3, DU145, and LNCaP used in this study were purchased from ATCC. LNCaP cells were grown in RPMI-1640 (Corning # 10-040-CV). PC-3 cells were cultured in DMEM/F12 (Corning # 10-092-CV). DU145 cells were grown in Dulbecco’s Modified Eagle Medium (DMEM, Corning #10-017-CV). The cells were maintained at a constant temperature of 37°C in a humidified incubator (5% CO2, / 95% air). Media was supplemented with 10% heat-inactivated fetal bovine serum (FBS) (# 10437028, Gibco), 1% penicillin-streptomycin (15070-063, Gibco).

### Establishment of Enzalutamide-resistant cells

Enzalutamide-resistant (ENZ-R) prostate cancer cell lines were generated from the AR-positive LNCaP and C4-2 parental cells through chronic exposure to progressively increasing concentrations of enzalutamide over six months. Initially, LNCaP and C4-2 cells were cultured in RPMI-1640 medium containing enzalutamide at a sub-lethal concentration (1μM). At approximately 90% confluence, the cells were subcultured and continuously exposed to gradually increasing concentrations of enzalutamide, up to 5μM. Cells that subsequently exhibited sustained exponential growth in the presence of 5μM enzalutamide were considered resistant and designated as enzalutamide-resistant (ENZ-R) cell lines. LNCaP ENZ-R and C4-2 ENZ-R were further maintained in the same media (RPMI 1640 media containing 5μM ENZ).

### Cell proliferation assay

Cells were seeded in a 96-well plate at a density of ∼ 3000 cells/well for 24 hours. After 24 hours of duration, cells were treated with varying concentrations of CA3 for up to 96 hours. Real-time proliferation was monitored using the Incucyte live-cell imaging system. Pictures were taken every 2 hours throughout the experiment. The data were used to calculate the cell proliferation rate (% confluency), and graphs were generated using the GraphPad Prism software.

### Colony formation assay

Colony formation assays were performed as described previously ^28^ . Cells were seeded in 12-well plates at ∼500 cells per well in growth media supplemented with vehicle or CA3 (0.5-0.75μM). After the indicated incubation period, cells were fixed and stained with 0.4% crystal violet for 30 minutes. After staining, wells were gently washed with distilled water and left to dry. Wells were imaged, and images were quantified in ImageJ to count the number of colonies.

### Transwell Invasion and Wound-Healing Migration Assays

Invasion assays were performed as described previously ^29^ , Briefly, the upper chambers of 24-well transwell inserts (8.0 μm pore size) were pre-coated with growth factor-reduced Matrigel (Matrigel (354234) at a 1:25 dilution in serum-free medium. Approximately 2 * 10^5 cells suspended in 300 μL of serum-free medium (containing specified concentrations of CA3) were seeded into the upper chamber. The lower chamber was filled with 10% FBS-supplemented medium to serve as a chemoattractant. Following the indicated incubation periods, non-invading cells on the upper surface of the membrane were removed with a cotton swab. Invaded cells on the lower surface were fixed, stained with 0.4% crystal violet, and photographed. Invaded cell populations were quantified across multiple random fields using ImageJ software.

For wound-healing migration assays, described previously ^30^ , cells were grown to a confluent monolayer in 6-well plates. A precise, linear wound track was introduced across the monolayer using a sterile 200 μL pipette tip. The wells were gently rinsed with PBS to remove cellular debris, and fresh media containing vehicle or CA3 (0.5–0.75 μM) were added. Micrographs of the wounded zones were captured at 0, 24, 48, and 72 h post-scratching. The percentage of wound closure was determined via ImageJ using the following formula: % Wound Closure = (Initial Area (0 h) - Residual Area (t h)) / Initial Area (0 h) * 100.

### RNA extraction and qPCR

RNA was extracted from the cells using the RNeasy Mini Kit (QIAGEN, 74106) according to the manufacturer’s instructions. We determined RNA concentration and purity using a NanoDrop spectrophotometer. RNA was reverse transcribed to complementary DNA using the iScript cDNA Synthesis Kit (Bio-Rad #1708891). Target cDNA was amplified using iTaq Universal SYBR Green Supermix (Bio-Rad, 172-5121) on a LightCycler 480 system (Roche) according to previously established methods ^31^ . The primers used in this study are provided in the table in the supplementary section. We used the delta-delta Ct method and normalized values to the housekeeping gene.

### Protein extraction and immunoblot

Protein extraction was performed as described previously ^32^ . Briefly, cells were lysed in RIPA buffer containing 150 mM NaCl, 10 mM Tris-HCl (pH 8.0), 1% Triton X-100, 5 mM EDTA, 0.1% SDS, 1% sodium deoxycholate, 1mM sodium orthovanadate, and a protease inhibitor cocktail (A32965, Pierce). After incubating the cells in the RIPA buffer for 30 minutes on ice, the cell lysates were sonicated with only 1-2 zaps. The cell lysate was then centrifuged at 14,000 rpm for 15 minutes at 4 °C. After centrifugation, the supernatant was collected and stored at −80 °C. Protein concentration in the samples was measured using the BCA Protein Assay Kit (23227). Protein samples were electrophoresed on 8-10% polyacrylamide gels or gradient gels (4-15% sodium dodecyl sulfate-polyacrylamide gel electrophoresis (SDS-PAGE; Bio-Rad, 4561084). Proteins were transferred to 0.45 μm polyvinylidene fluoride (PVDF) or nitrocellulose membranes, and non-fat dry milk was used to block non-specific binding, followed by incubation with primary and HRP-coupled secondary antibodies to identify the proteins of interest. The chemiluminescent ECL solution (Immobilon® Forte #WBLUF0500, Millipore Sigma) was used to visualize the blots via the Versa Doc imaging system or Chemidoc imaging system (Bio-Rad).

#### Immunofluorescence and Quantitation

Immunofluorescence assays were performed as described previously^29^ . Briefly, cells were cultured on 8-chambered glass slides, fixed with 4% formaldehyde for 30 min at room temperature, and permeabilized with an ice-cold 20% methanol/80% acetone mixture for 5 min on ice. Non-specific binding sites were blocked with 5% bovine serum albumin (BSA) in PBS for 1 h. Cells were incubated at room temperature with primary antibodies against YAP-11 or γ-H2AX for 1 h, followed by fluorophore-conjugated secondary antibodies for 1 h. Nuclei were counterstained with DAPI. Slides were mounted using fluorescence mounting media and imaged on an epifluorescence microscope. Fluorescence intensity was quantified across multiple fields using ImageJ.

### Xenograft mouse model

Experiments were conducted with approval from the Institutional Animal Care and Use Committee at Louisiana State University Health Science Center, New Orleans. Six-week-old nude male mice were purchased from The Jackson Laboratory, USA, and acclimatized for one week before the experiment. Animals had ad libitum access to food and water and were housed in cages with controlled temperature and humidity. Before injection, mice were anesthetized with isoflurane. PCa cells (DU145 and PC3; 1 × 10⁶) were suspended in Matrigel-containing media at a 1:1 ratio and injected subcutaneously into each flank of the right and left limbs using a 28G needle. Mice were monitored until they regained full consciousness. CA3 stock solution was prepared in DMSO and further diluted in PBS to the working concentration. Once tumors were palpable, mice were randomly assigned to treatment or control groups. Mice in the treatment group received CA3 intraperitoneally at 1.5 mg/kg body weight (100 μL) on alternate days until the endpoint. Control mice received 100 μL of vehicle (PBS) in the same manner. Body weight and tumor volume were measured twice weekly until the study endpoint. Tumor volume was calculated as 0.5 × (L × W²). At the end of the experiment, mice were sacrificed, xenografts were excised, tumor tissues were weighed, and samples were saved for further analysis.

#### H & E staining and Immunohistochemistry

H&E staining was performed on paraffin-embedded tissues using the protocol described previously ^33^ . IHC was performed using the protocol described previously ^32^ . Chromogenic detection was performed using the manufacturer’s protocol for VECTASTAIN reagent (SK-4105; Vector Labs). We counted Ki67-, PCNA-, and γ-H2AX-positive cells in 5 fields at 20× magnification (ImageJ software), and we quantified and analyzed the data using statistical software.

### Masson’s Trichrome Staining

We evaluated collagen fibers in paraffin-embedded tissue sections using a commercial Trichrome Stain Kit (Abcam #150686). Briefly, sections were deparaffinized, rehydrated, and incubated in Bouin’s fluid as a mordant. Nuclei were stained with Weigert’s Iron Hematoxylin, and the cytoplasm was subsequently stained with Biebrich Scarlet/Acid Fuchsin solution. The slides were differentiated in a phosphomolybdic/phosphotungstic acid solution and immediately counterstained with Aniline Blue to target collagen fibers. Finally, the sections were treated with an acetic acid solution, dehydrated, cleared, and mounted for imaging.

### TUNEL Assay

To evaluate apoptosis, a Terminal deoxynucleotidyl transferase dUTP nick end labeling (TUNEL) assay was performed using a commercial detection kit (#25879, Cell Signaling Technology) according to the manufacturer’s instructions. Briefly, tissue sections were deparaffinized in xylene and rehydrated through a graded series of ethanol concentrations. Antigen retrieval was conducted in a 10 mM sodium citrate buffer for two 5-minute cycles. The slides were then incubated in TUNEL equilibration buffer for 5 minutes, followed by incubation with the TUNEL reaction mixture for 2 hours at 37°C in the dark. After washing with PBST for 5 minutes, the sections were counterstained with DAPI for 5 minutes at room temperature. Finally, the slides were mounted, and fluorescent images were captured for analysis. TUNEL-positive cells were quantified using ImageJ software (NIH, Bethesda, MD, USA).

### Annexin V staining

Cell death was analyzed using the Annexin V-FITC Early Apoptosis Detection Kit (#6592). Briefly, cells cultured under the respective experimental conditions were treated with the drug, then harvested by trypsinization and washed with PBS after incubation. Cells were subsequently stained with Annexin V-FITC and PI according to the kit protocol. Data were acquired using a flow cytometer and analyzed with FlowJo software. Graphs were generated in GraphPad Prism. In this assay, cells positive for Annexin V and negative for PI were considered early apoptotic, while cells positive for both Annexin V and PI were considered late apoptotic.

### Analysis of Clinical Data

The normalized gene expression, as well as phenotypes including overall survival time, in the Metastatic Prostate Adenocarcinoma data set (MCTP, Nature 2012) and gene expression across prostate cancer cell lines (Next Generation Characterization of the Cell Line Encyclopedia) were downloaded from cBioPortal (https://www.cbioportal.org/) ^34,35^ . The data were analyzed in GraphPad Prism.

### Statistical Analysis

Statistical analysis was performed using GraphPad Prism. Normality tests were conducted on all immunoblot assay data to assess whether the data were normally distributed, thereby allowing the use of parametric tests. Two-group comparisons were analyzed using a two-tailed Student’s t-test. Quantitative data were expressed as mean ± standard error of the mean (SEM). Statistical significance was defined as p<0.05. All experiments were performed in triplicate with at least two independent biological replicates.

## Results

### 1. YAP-1/TAZ-TEAD transcriptional activity is enriched in enzalutamide-resistant PCa cells

To test whether enzalutamide resistance is functionally linked to Hippo-YAP-1/TAZ activation, we profiled the transcriptional and translational status of key pathway components in isogenic enzalutamide-resistant models.

Quantitative RT-PCR analysis showed substantial up-regulation of multiple canonical Hippo/TEAD target genes, specifically COL4A3, FGF2, ITGB2, DLC1, TOP2A, AMOTL2, and ETV5 in LNCaP ENZ-R cells relative to their parental counterparts (Fig. 1A). To test whether these gene alterations correlate with increased protein expression, we examined total protein levels by Western blot (Fig. 1B, C). Notably, both LNCaP ENZ-R and C4-2 ENZ-R cells exhibited marked increases in total protein levels of YAP-1, TAZ, and TEAD compared with parental controls (Fig. 1D, E). Because nuclear translocation is required for YAP-1/TAZ to act as transcriptional co-activators, we performed high-resolution immunofluorescence imaging to assess subcellular distribution. Compared with parental cells, enzalutamide-resistant variants displayed a pronounced shift toward nuclear YAP-1 localization, validating increased transcriptional activation of the axis (Fig. 1F, G). Taken together, these data establish that enzalutamide resistance in prostate cancer cells is accompanied by considerable activation and nuclear enrichment of the Hippo-YAP-1/TAZ signaling cascade.

**Figure 1:**
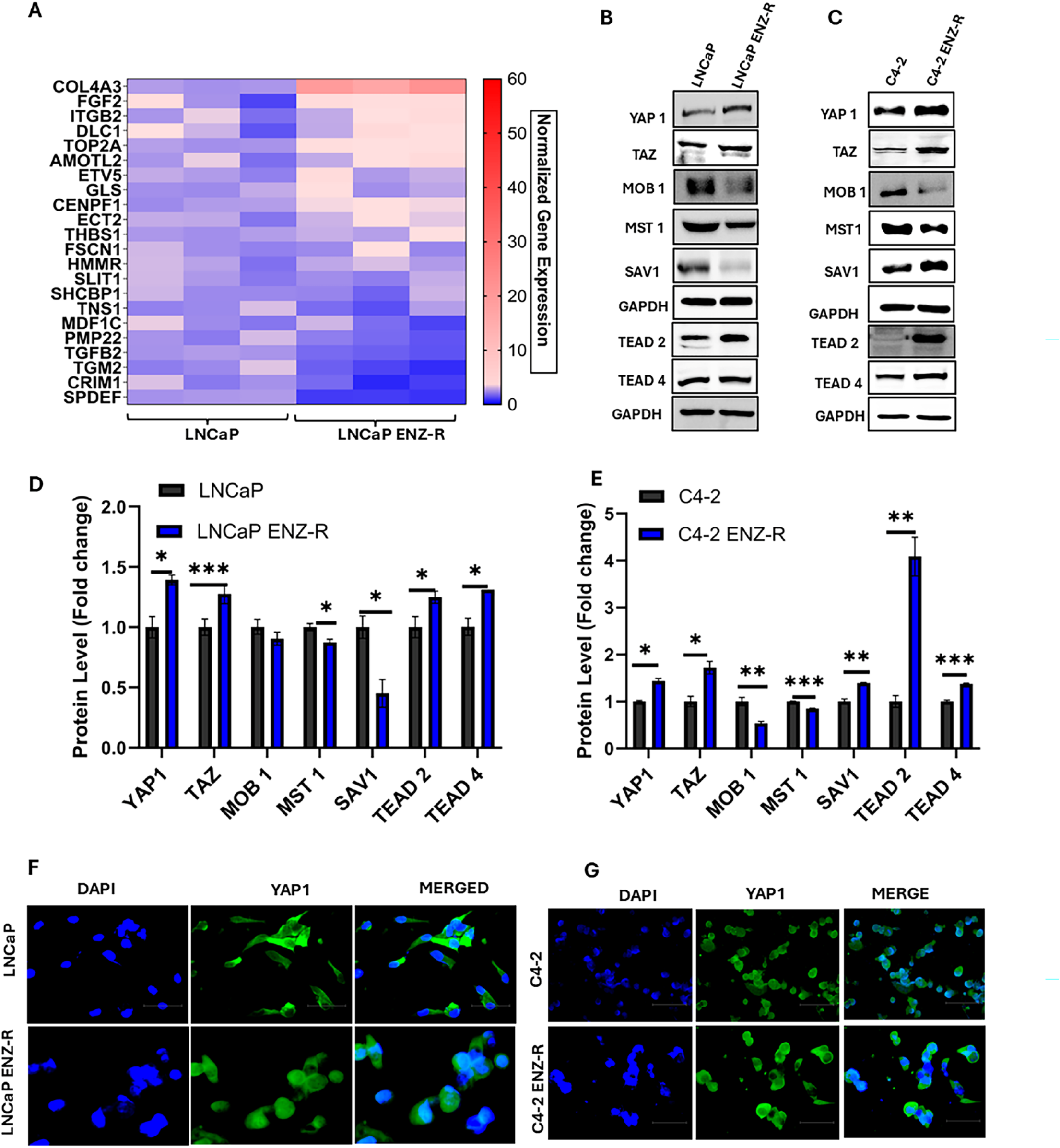
YAP-TAZ-TEAD transcriptional activity is enriched in Enzalutamide-resistant PCa: Expression of Hippo YAP/TAZ-TEAD transcriptional targets in LNCaP and LNCaP ENZ-R cells (A); Western blots of Hippo YAP/TAZ-TEAD signaling pathway proteins in LNCaP (B); C4-2 (C); and their respective ENZ-R derivative cell lines. Quantitative analysis of Hippo YAP/TAZ-TEAD protein blots from LNCaP/LNCaP ENZ-R cells (D) and C4-2/C4-2 ENZ-R cells (E); data are presented as mean ± S.D. of 3 experiments, * p≤0.05, ** p≤0.01, and *** p≤0.001 compared to respective controls. Immunofluorescence images (IMF) of YAP in LNCaP/LNCaP ENZ-R (F) and C4-2/C4-2 ENZ-R cells (G).

### 2. Pharmacological disruption of YAP-1/TAZ-TEAD suppresses proliferation, migration, and clonogenicity across diverse PCa models

To evaluate the functional requirement of the YAP-1/TAZ-TEAD axis, we tested the therapeutic effects of CA3, a potent small-molecule inhibitor of the physical interaction between YAP-1/TAZ and TEAD. Androgen-sensitive (LNCaP) and androgen-independent (DU145, PC3) cell lines were exposed to gradient doses of CA3 (0, 0.5, 0.75, and 1.0 μM), and cell proliferation kinetics were monitored in real time using the IncuCyte system. CA3 treatment triggered robust, concentration- and time-dependent reductions in cell confluence across all three lines (Fig. 2A, B, C). CA3 strongly inhibited growth at 0.5 μM, with near-complete arrest at 0.75 μM and 1.0 μM, highlighting CA3’s broad efficacy regardless of baseline AR status. We next evaluated whether CA3 impairs PCa cell migration using a scratch wound-healing assay.

**Figure 2:**
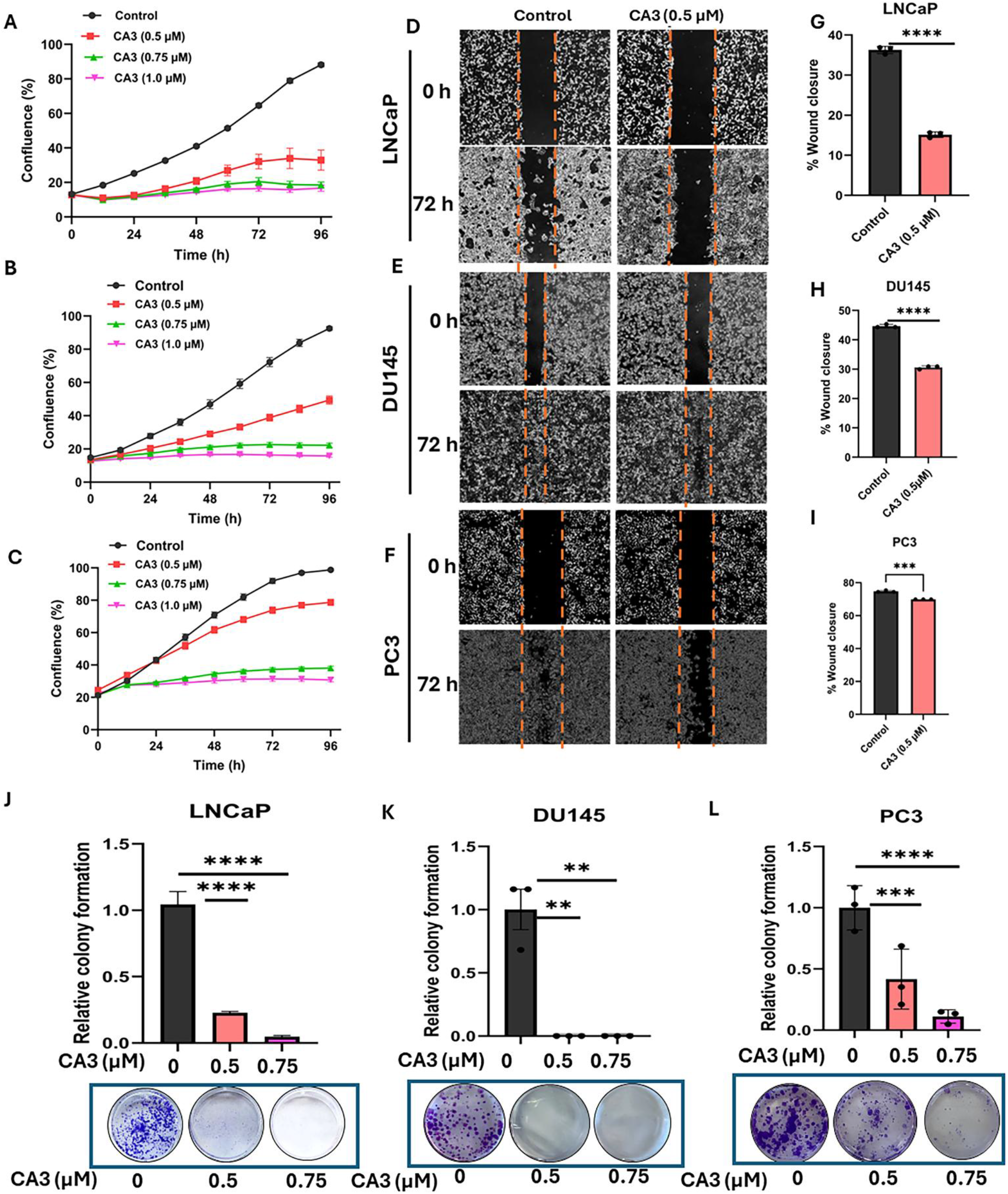
YAP/TAZ-TEAD activity is essential for PCa cells. Cell growth in LNCaP (A), DU145 (B), and PC3 (C) cells after treatment with increasing concentrations of CA3 (YAP/TAZ-TEAD inhibitor). Representative images of the scratch wound assay in LNCaP (D), DU145 (E), and PC3 (F) cells after treatment with CA3. Quantification of these data (n=3) using ImageJ is presented graphically on the right side (G, H, I); colony formation assay quantification and representative images are shown at the bottom in LNCaP (J), DU145 (K), and PC3 (L). Error bars indicate the standard deviation of experiments performed in triplicate, and significance is shown as * p≤0.05, ** p≤0.01, *** p≤0.001, and **** p≤0.0001 compared to respective controls using a Student t-test.

Treatment with 0.5μM or 0.75μM CA3 significantly delayed wound closure over a 72-h tracking period in all cell lines compared to controls (Fig. 2D, E, F). Quantitative analysis confirmed a marked reduction in final wound-closure percentage, indicating that active YAP-1/TAZ-TEAD signaling is required for prostate cancer cell motility (Fig. 2G, H, I). Furthermore, long-term colony formation assays in all three cell lines showed that low-dose CA3 (0.5 and 0.75μM) drastically reduced both the total number and average colony size over 10 days, indicating impaired clonogenic survival (Fig. 2J, K, L).

### 3. YAP-1/TAZ-TEAD Inhibition suppresses proliferation and aggressive phenotypes specifically in enzalutamide-resistant LNCaP cells

Because enzalutamide-resistant cells enrich for YAP-1/TAZ signaling, we tested whether targeting this pathway could exploit a key therapeutic vulnerability in resistant populations. We treated LNCaP ENZ-R cells with enzalutamide, CA3, and both in combination (Fig. 3). LNCaP ENZ-R cells grew rapidly despite continuous exposure to 5μM enzalutamide, confirming their resistant phenotype. In contrast, CA3 treatment bypassed this resistance, inducing dose-dependent growth suppression in the ENZ-R cell lines, with maximum efficacy at 0.75μM and 1.0μM (Fig. 3A).

**Figure 3:**
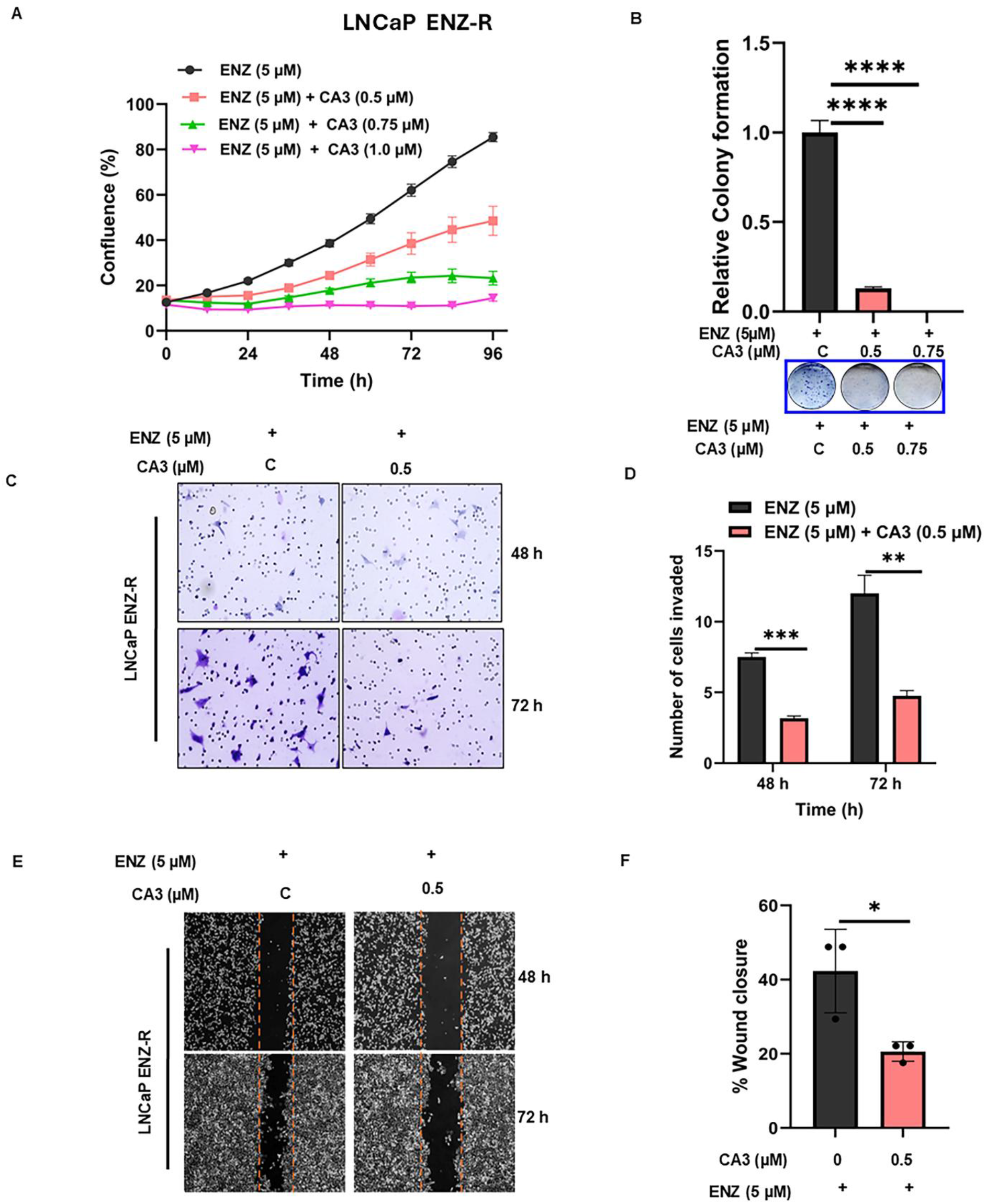
Pharmacological inhibition of TEAD activity sensitizes LNCaP ENZ-R cells. Cell growth inhibition rate curve for LNCaP ENZ-R cells (A), showing decreased cell proliferation upon treatment with CA3 alone and in combination with ENZ. Representative images of colony formation in LNCaP ENZ-R cells (B) and their respective quantification. Data are presented as Mean ± SD (n=3; significance is shown as **** p≤0.0001 compared with the respective control). Representative microscopic images of the invasion assay in LNCaP ENZ-R cells and their quantification using ImageJ software in the bottom panel (C); scratch assay for cell migration in LNCaP ENZ-R cells treated with CA3 and ENZ (D). Images were taken at 0, 48, and 72 h after drug treatment. Quantitative data are Mean ± SD (n=3, * p≤0.05, ** p≤0.01, and *** p≤0.001 compared to respective controls).

To confirm that this therapeutic vulnerability shows a broad mechanistic dependency on functional Hippo transcription rather than a compound-specific artifact of CA3, we evaluated two additional structurally distinct small-molecule YAP-1/TAZ-TEAD inhibitors, GNE-7883 and K-975, in parallel live-cell growth assays. Consistent with our primary CA3 findings, both GNE-7883 and K-975 induced robust, concentration-dependent suppression of cell growth across both parental and enzalutamide-resistant cell states (data not shown). Moreover, siRNA-mediated knockdown of YAP-11 and TAZ decreased cell proliferation and colony formation in LNCaP ENZ-R cells, and CERES dependency score maps for WWTR1 also confirm that WWTR1 is indispensable for most of the prostate cancer cell lines (Fig. S1). This validation strongly supports a true therapeutic class effect following disruption of the nuclear YAP-1/TAZ-TEAD complex.

This trend was further reflected in long-term clonogenic survival assays. LNCaP ENZR cells retained robust colony-forming capability under 5μM enzalutamide (Fig. 3B). Treatment with 0. 5μM or 0. 75μM CA3 significantly reduced colony formation in these resistant populations (****p< 0. 0001), leading to nearly complete eradication at 0. 75μM (Fig.4B). Additionally, CA3 exposure significantly decreased transwell invasion at both 48 h and 72 h compared to enzalutamide alone (**p< 0. 001 and **p< 0.01, respectively; Fig. 3C, D). Wound-healing migration rates were also lower, with CA3-treated cells achieving only ∼20% wound closure at 72 h compared to ∼40% in the enzalutamide-treated ENZR controls (*p< 0.05; Fig. 3E, F).

**Figure 4:**
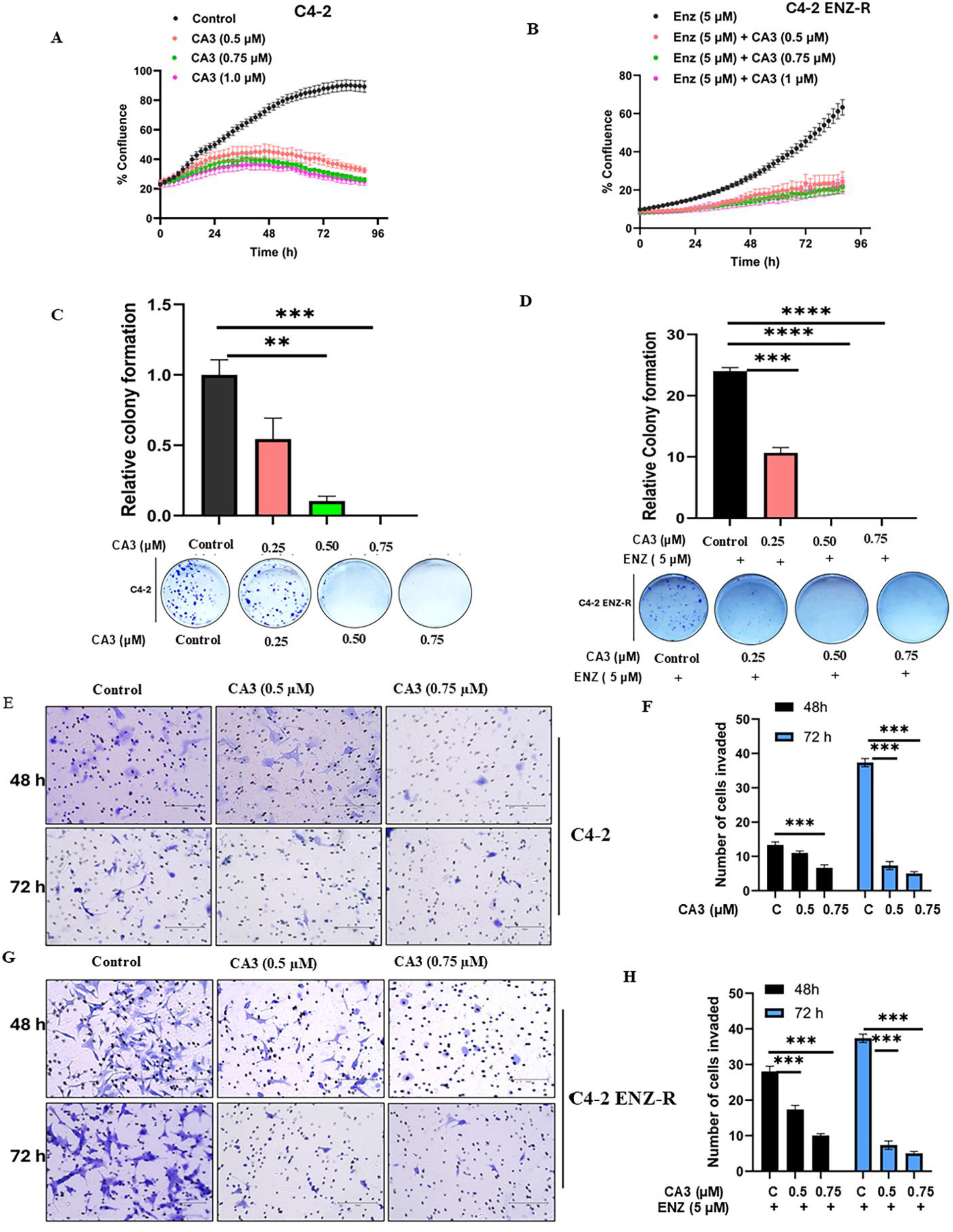
Pharmacological Inhibition of TEAD Activity sensitizes C4-2 Prostate Cancer Cells. Growth response curves in C4-2 cells (A) and C4-2 ENZ-R cells (B) after drug treatment, monitored over 96 h in Incucyte. Graphs showing relative colony formation in C4-2 cells (C) and C4-2 ENZ-R cells (D). At the bottom of the graphs are microscopic representations of colonies formed in the aforementioned cell lines. Cell invasion images in C4-2 (E) and C4-2 ENZ-R cells (G) using a Boyden chamber, with quantifications of the images on the right (F, H) (n=3). Error bars indicate the standard deviation of experiments performed in triplicate, and significance is shown as * p≤0.05, ** p≤0.01, *** p≤0.001, and **** p≤0.0001 compared to respective controls using a Student’s t-test.

### 4. CA3 suppresses proliferation, clonogenicity, and transwell invasion in parental and enzalutamide-resistant C4-2 cells

To confirm the generalizability of these findings in an independent advanced PCa context, we evaluated CA3 in the C4-2 progression model (Fig. 4). CA3 administration markedly reduced the proliferation rate of parental C4-2 cells across a concentration range of 0.5–1.0μM (Fig. 4A). Similarly, isogenic C4-2 ENZ-R cell lines proliferated continuously in media containing 5μM enzalutamide, but addition of CA3 (0.5–1.0μM) reduced cell proliferation confirming that CA3 retains anti-proliferative efficacy in enzalutamide-resistant cell states (Fig. 4B). Long-term colony assays showed a similar, dose-dependent decrease in clonogenic potential for both parental C4-2 and C4-2 ENZ-R variants. In parental C4-2 cells, colony growth was severely restricted at 0.5μM and 0.75μM CA3 (**p< 0.01 and ***p< 0.001; Fig. 5C). C4-2 ENZ-R cells sustained strong colony growth under enzalutamide monotherapy, but addition of 0.5μM or 0.75μM CA3 led to near-complete suppression (**p< 0.001 and ****p< 0.0001, respectively; Fig. 4D). Furthermore, transwell assays confirmed that CA3 treatment significantly reduced cell invasion through Matrigel at 48 h and 72 h in both the parental line (Fig. 4E, F) and the enzalutamide-resistant line compared with enzalutamide alone (***p< 0.001; Fig. 4G, H).

**Figure 5:**
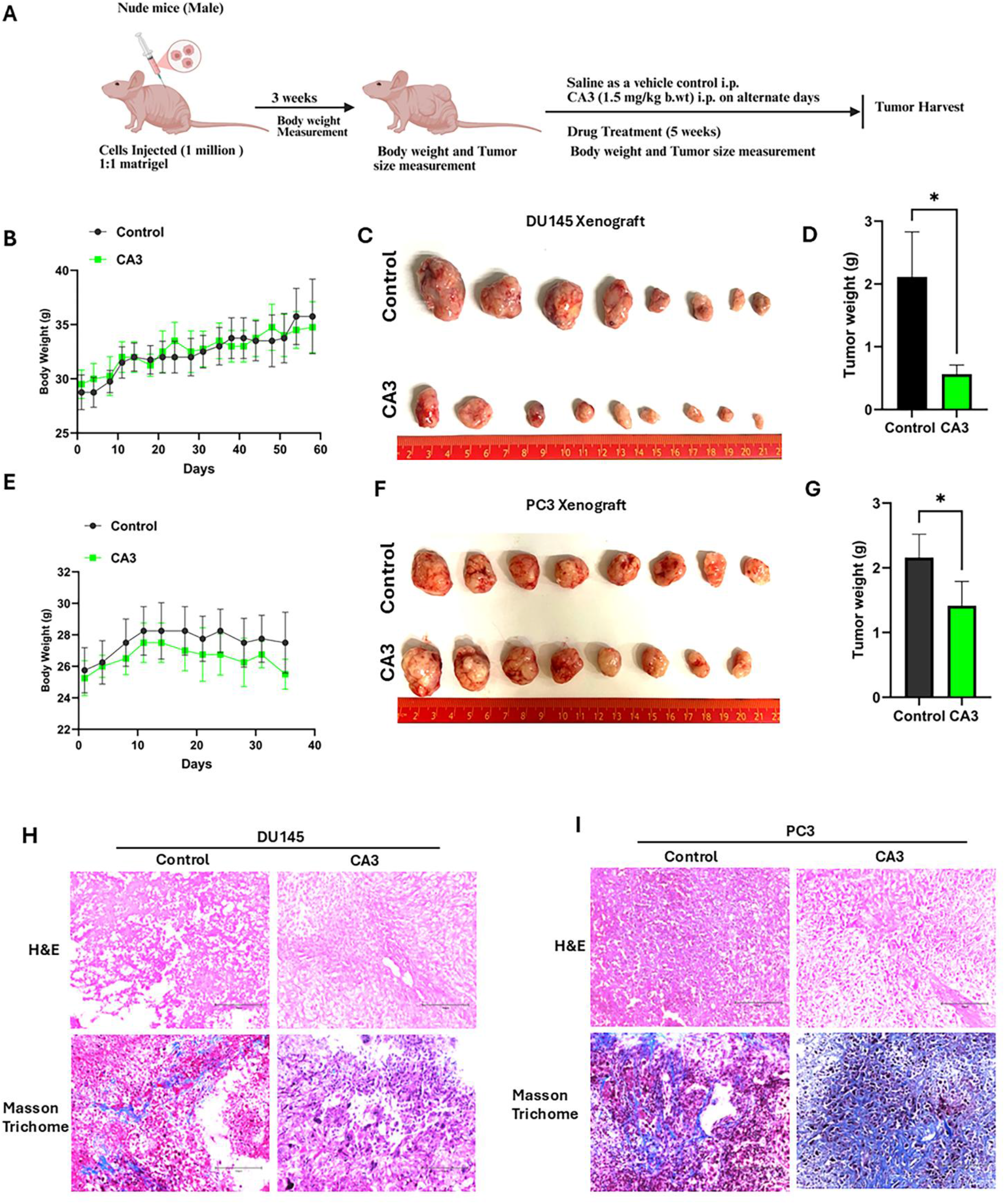
CA3 (YAP/TAZ-TEAD inhibitor) inhibits tumor growth in PCa xenografts in vivo. Schematic representation of the dosing regimen in tumor-bearing mice (A); graph showing body weight of DU145 (B) and PC3 (E) tumor-bearing mice. Body weights were recorded twice a week, and error bars in the graph represent Mean ± SD. Representative images of tumors from mice injected with vehicle and CA3 (1.5 mg/kg b.wt) in DU145 xenograft (C) and PC3 xenograft mice (F), respectively. The right-hand side panel depicts tumor weights from CA3-treated mice and their respective controls (D, G). Representative H&E and Masson trichrome staining images from DU145 (H) and PC3 (I) tumors excised from mice treated with CA3 and their controls.

### 5. Systemic CA3 administration suppresses tumor growth and induces histopathological remodeling *in vivo*

To translate our *in vitro* findings into a preclinical animal model, we evaluated CA3’s therapeutic efficacy using DU145 and PC3 subcutaneous xenografts in male nude mice. Once tumors were established, CA3 (1.5 mg/kg body weight, i.p.) was administered on alternate days for 5 weeks (Fig. 5A). Systemic CA3 administration was well tolerated; no adverse behaviors or significant changes in total body weight were observed among control and treated cohorts throughout the study period (Fig. 5B, E). In the DU145 xenograft model, CA3 treatment reduced tumor burden (Fig. 5 C), resulting in a significant decrease in final tumor weight relative to vehicle controls (*p< 0.05; Fig. 5D). A similar anti-tumor response was observed in the PC3 xenograft cohort (Fig. 5 F), with treated mice showing significantly lower average tumor weights than their control counterparts (*p< 0.05; Fig. 5G). H&E staining revealed noticeable alterations in tissue architecture and cellular morphology in CA3-treated tumors from both models (Fig. 5H, I). Furthermore, Masson’s Trichrome staining demonstrated treatment-associated stromal remodeling. CA3-treated tumors showed increased collagen deposition (blue staining), particularly in the PC3 cohort, indicating significant architectural remodeling of the tumor microenvironment following treatment (Fig. 5H, I).

### 6. CA3 decreased expression of PCNA and KI-67, markers of cell proliferation in Prostate Cancer Xenografts in vivo

To clarify the cellular mechanisms underlying the observed reduction in tumor burden *in vivo*, we evaluated treated xenograft tissue sections for changes in cell proliferation and survival. Immunohistochemical staining for the established proliferation markers PCNA and Ki67 displayed a marked decrease in proliferative indices following CA3 treatment (Fig. 6A, B). In DU145 tumor sections, CA3 significantly decreased the percentage of PCNA-positive cells (***p< 0.001) and Ki67-positive cells (*p<0.05; Fig. 6B) compared with control sections. Consistent with this, PC3 xenograft tissues from the CA3 cohort exhibited lower expression of both PCNA (*p< 0.05) and Ki67 (***p< 0.001; Fig. 6B). To test whether the loss of tumor cell mass was also driven by programmed cell death, we performed TUNEL staining on the xenograft tissue sections (Fig. 6C, D).

**Figure 6:**
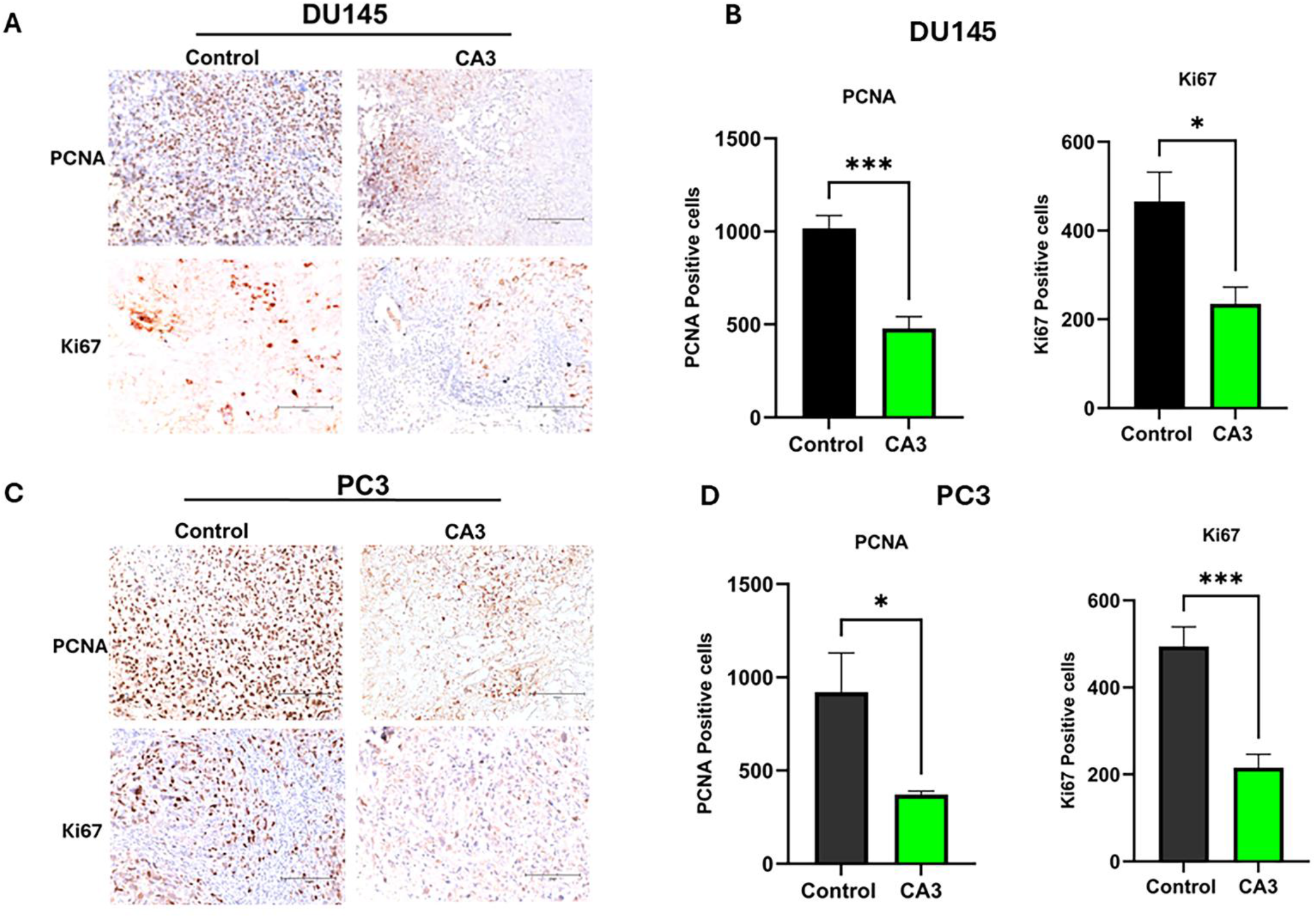
YAP/TAZ TEAD inhibitor decreases tumor growth and proliferation in PCa xenografts. Representative immunohistochemistry (IHC) photomicrographs showing PCNA and Ki67 proliferation markers in control and CA3-treated tumors from DU145 (A) and PC3 (C) xenografts. On the right side is the quantification of these data (B, D). Quantitative data are presented as Mean ± SD (n=3, * p≤0.05 and *** p≤0.001 compared to respective controls).

### 7. YAP-1/TAZ TEAD inhibitor induces apoptosis *in vitro* and *in vivo*

To confirm the mechanism of cell death induced by the CA3 inhibitor, we performed Annexin V/PI dual staining in vitro. Flow cytometric analysis demonstrated a dose-dependent increase in both early and late apoptotic populations after CA3 treatment (0.5 and 0.75 μM) in DU145 and PC3 cells (Fig. 7A, B). To confirm these results in vivo, we performed TUNEL assays on tissue sections harvested from tumor xenografts. DU145 tumors from the CA3-treated cohort displayed a marked increase in the abundance of fluorescent TUNEL-positive apoptotic cells compared with vehicle-treated control sections, as confirmed by quantitative analysis (**p < 0.01; Fig. 7E, F). Similarly, treated PC3 tumors exhibited a significant increase in the TUNEL-positive apoptotic index (*p < 0.05; Fig. 7G, H). Collectively, the results show that CA3’s in vivo anti-tumor efficacy is driven by lowered proliferation and accelerated apoptosis.

**Figure 7:**
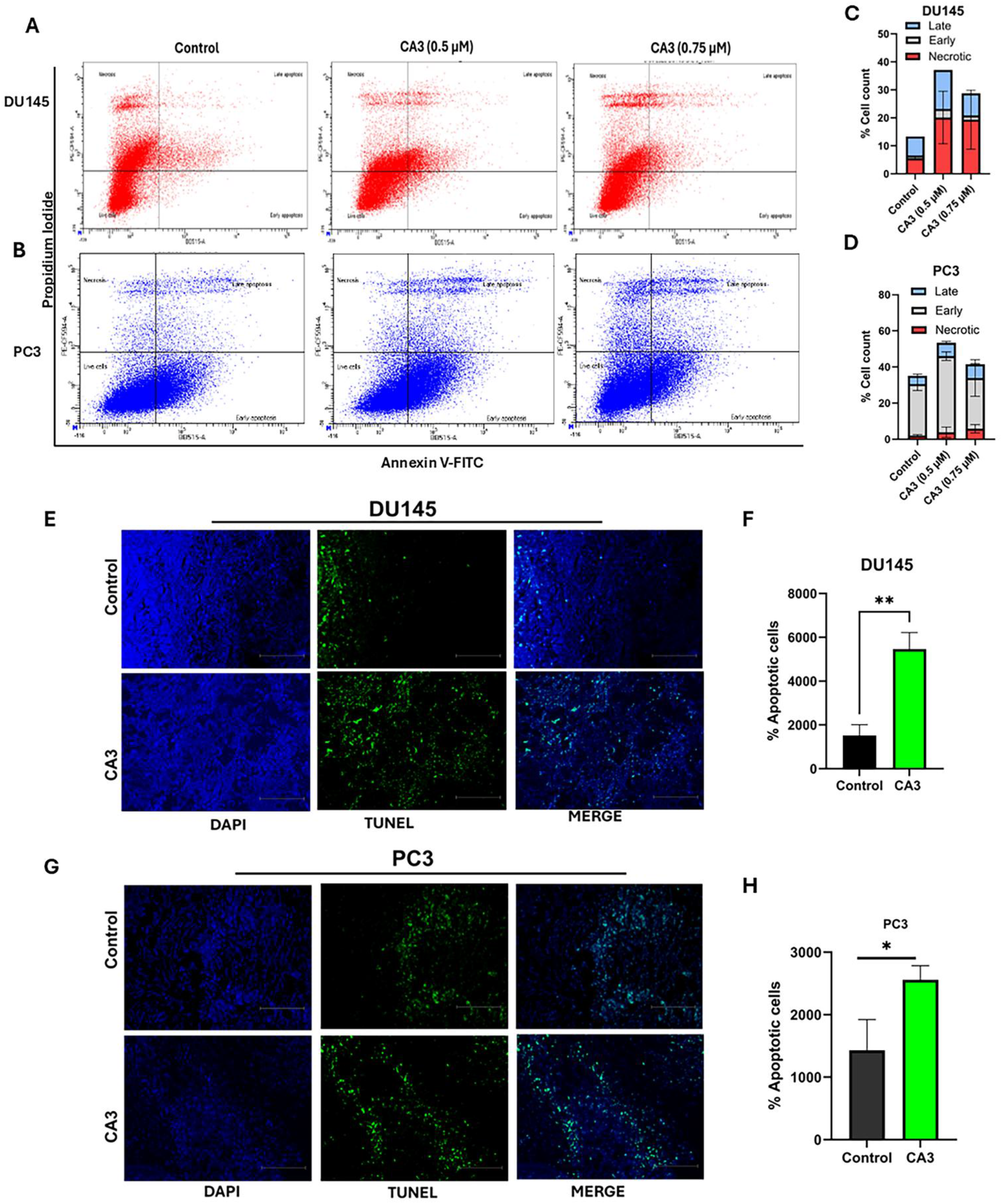
YAP/TAZ TEAD inhibitor induces apoptosis in vitro and in vivo. Untreated and CA3-treated PCa cells were collected and stained with Annexin V-FITC and PI. DU145 (A) and PC3 (B) are representative dot plots for Annexin V and PI staining. Histograms represent the percentage of cells undergoing early apoptosis, late apoptosis, and necrosis as determined by Annexin V/PI staining (C, D). Experiments were performed independently three times. Detection and quantification of apoptosis (TUNEL assay) in DU145 (E, F) and PC3 (G, H) xenografts, respectively. The right side represents quantification of these data. Data are presented as Mean ± SD (n=3; * p≤0.05 and ** p≤0.01 compared to respective controls).

### 8. CA3 triggers pronounced DNA damage accumulation and impairs homologous recombination repair

To test whether CA3’s anti-proliferative and pro-apoptotic activities are linked to genotoxic stress, we evaluated γ-H2AX accumulation, a well-established marker of DNA double-strand breaks. DU145 and PC3 cells were treated with 0.5 μM CA3 and analyzed by immunofluorescence microscopy (Fig. 8A, B). Vehicle-treated DU145 cells showed minimal baseline γ-H2AX signals, whereas CA3 treatment induced intense, nuclear-localized γ-H2AX foci. Quantification of fluorescence intensity confirmed a significant increase in DNA damage post-treatment (**p < 0.01; Fig. 8C). An identical response was observed in the PC3 cell line (Fig. 8D). Although baseline γ-H2AX signals were detectable in untreated control PC3 cells, exposure to 0.5 μM CA3 led to a substantial increase in nuclear γ-H2AX fluorescence intensity (***p < 0.0001; Fig. 8E).

**Figure 8:**
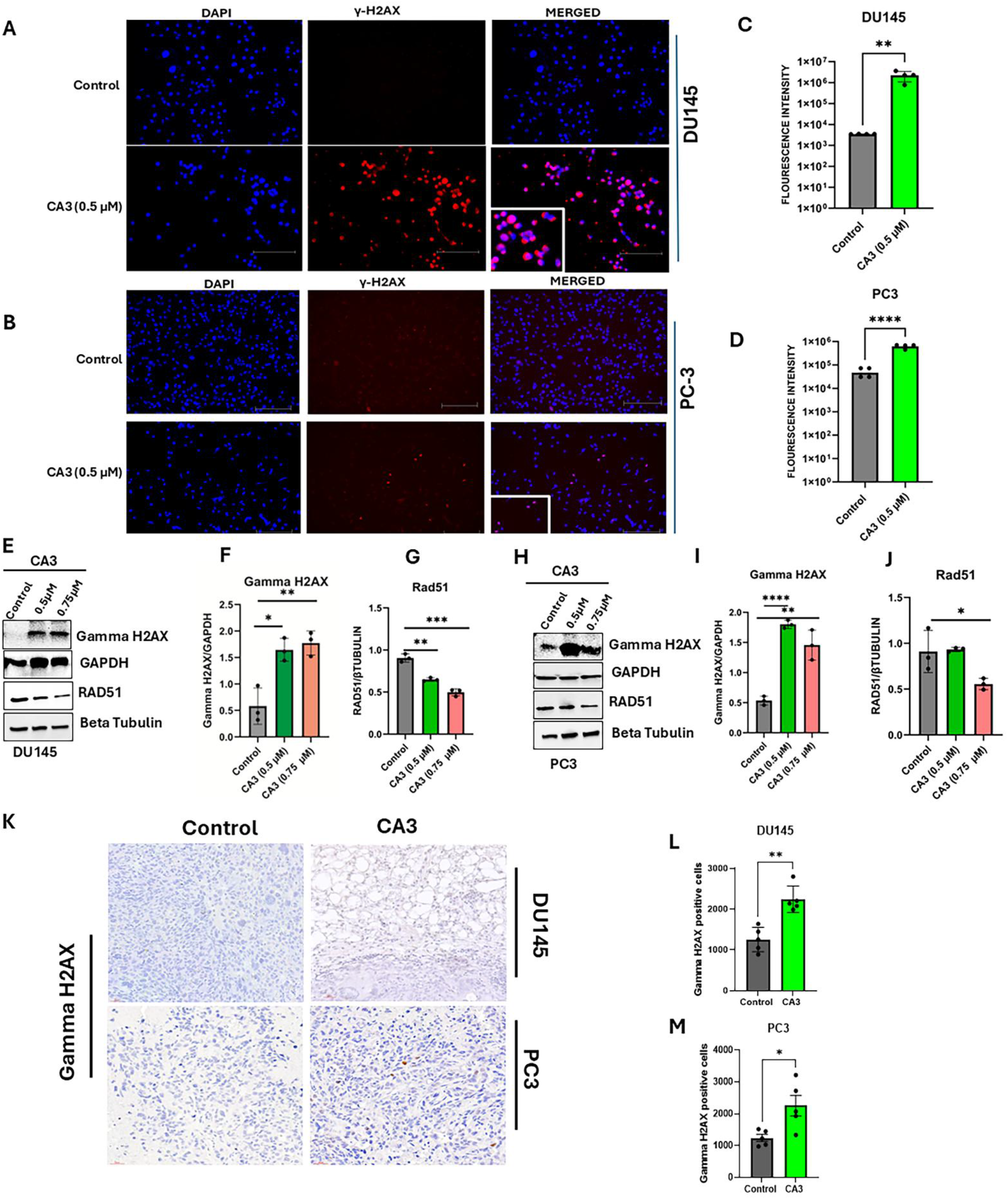
CA3 Treatment induces DNA damage in vitro and in vivo. Representative images of γ-H2AX staining in DU145 (A) and PC3 cells (B). On the right side is the quantification of these data (C, D). Error bars indicate the standard deviation of experiments performed in triplicate, and significance is shown as ** p≤0.01 and **** p≤0.0001 compared to the respective controls using a Student’s t-test. Western blot showing increased protein expression of γ-H2AX and Rad51 after CA3 treatment in DU145 (E) and PC3 cells (H). On the right is the quantification of the bands with respect to the control (F, G, I, J). Data are represented as Mean ± SD (n=3, * p≤0.05, ** p≤0.01, *** p≤0.001, and **** p≤0.0001 compared to the respective controls). Representative images of γ-H2AX staining in DU145 and PC3 xenograft tissues (K); on the left are the bar graphs showing an increase in γ-H2AX-positive cells (L, M); Error bars indicate the standard deviation of experiments performed in triplicate, and significance is shown as * p≤0.05 and ** p≤0.01 compared to the respective controls using a Student’s t-test.

To corroborate these data at the protein level, we evaluated γ-H2AX and Rad51 expression in DU145 and PC3 cells via Western blotting (Fig. 8E, H). Consistent with our immunofluorescence data, CA3 treatment significantly upregulated γ-H2AX protein expression in both cell lines (*p < 0.05, ***p < 0.001; Fig. 8F, I). Given the induction of double-strand breaks, we next investigated Rad51 expression, an essential part of the homologous recombination repair pathway (Fig. 8E, H). Quantitative analysis of the immunoblots showed a significant, dose-dependent decrease in Rad51 protein levels after treatment with 0.5 μM and 0.75 μM CA3 in both cell lines (Fig. 8G, J), indicating compromised DNA double-strand break repair. Notably, total Chk1 and its activated form, phospho-Chk1, remained unchanged (data not shown), suggesting that this treatment does not modulate classical DNA damage checkpoint signaling. To extend these data in vivo, we performed immunohistochemical analysis on DU145 and PC3 xenograft tumor tissues. CA3-treated tumors exhibited a marked increase in γ-H2AX staining intensity that closely mirrored our in vitro findings (Fig. 8K, L, M). Taken together, these data show that CA3-mediated pharmacological inhibition of the YAP1/TAZ-TEAD complex induces severe genotoxic stress by accelerating DNA double-strand break accumulation and impairing homologous recombination-mediated repair.

### 9. Elevated YAP1/WWTR1 associated with poor therapeutic response in advanced prostate cancer patients

We examined the association between WWTR1 (TAZ) and YAP-1 mRNA expression and clinical survival outcomes using a public human prostate cancer dataset (Metastatic Prostate Adenocarcinoma (MCTP, Nature 2012). We discovered that WWTR1 transcript levels were inversely correlated with overall patient survival, with higher expression tracking with shorter survival times (Spearman r = −0.40, P = 0.0272; Pearson r = −0.35, P = 0.0544; Fig. 9A). Negative trends were also observed between higher WWTR1 expression and patient survival following first-line hormone therapy (Spearman r = −0.24, P = 0.191; Pearson r = −0.27, P = 0.138; Fig. 9C) and first-line chemotherapy (Spearman r = −0.22, P = 0.241; Pearson r = −0.35, P = 0.0608; Fig. 9E). We observed similar trends between YAP-1 expression and patient outcomes (Fig. 9B, D, F), however, these latter trends did not reach statistical significance. We also examined YAP-1, WWTR1, and TEAD mRNA expression across a comprehensive panel of human PCa cell lines representing distinct stages and phenotypes. YAP-1 mRNA expression varied, with elevated levels in LNCaP, VCaP, PC3, DU145, and MDA-PCa-2b cell lines and low levels in NCI-H660 cells (Fig. 9 G). WWTR1 expression varied across the cell lines, with the highest transcripts in the aggressive, AR-independent cell lines DU145 and PC3 and the neuroendocrine model NCI-H660, whereas low levels were found in androgen-sensitive LNCaP, VCaP, and MDA-PCa-2b models (Fig. 9 H). Analysis of TEAD family members revealed heterogeneous expression profiles across the cell line panel (Fig. 9 I). Notably, TEAD1 and TEAD2 were highly expressed in aggressive cell lines, including DU145, PC3, VCaP, and NCI-H660, whereas the relative abundance of TEAD3 and TEAD4 varied across models. Taken together, these data show that YAP-1, WWTR1(Taz), and TEAD are clinically relevant targets, suggesting that WWTR1 is a potential driver of therapy resistance in aggressive prostate cancer.

**Figure 9:**
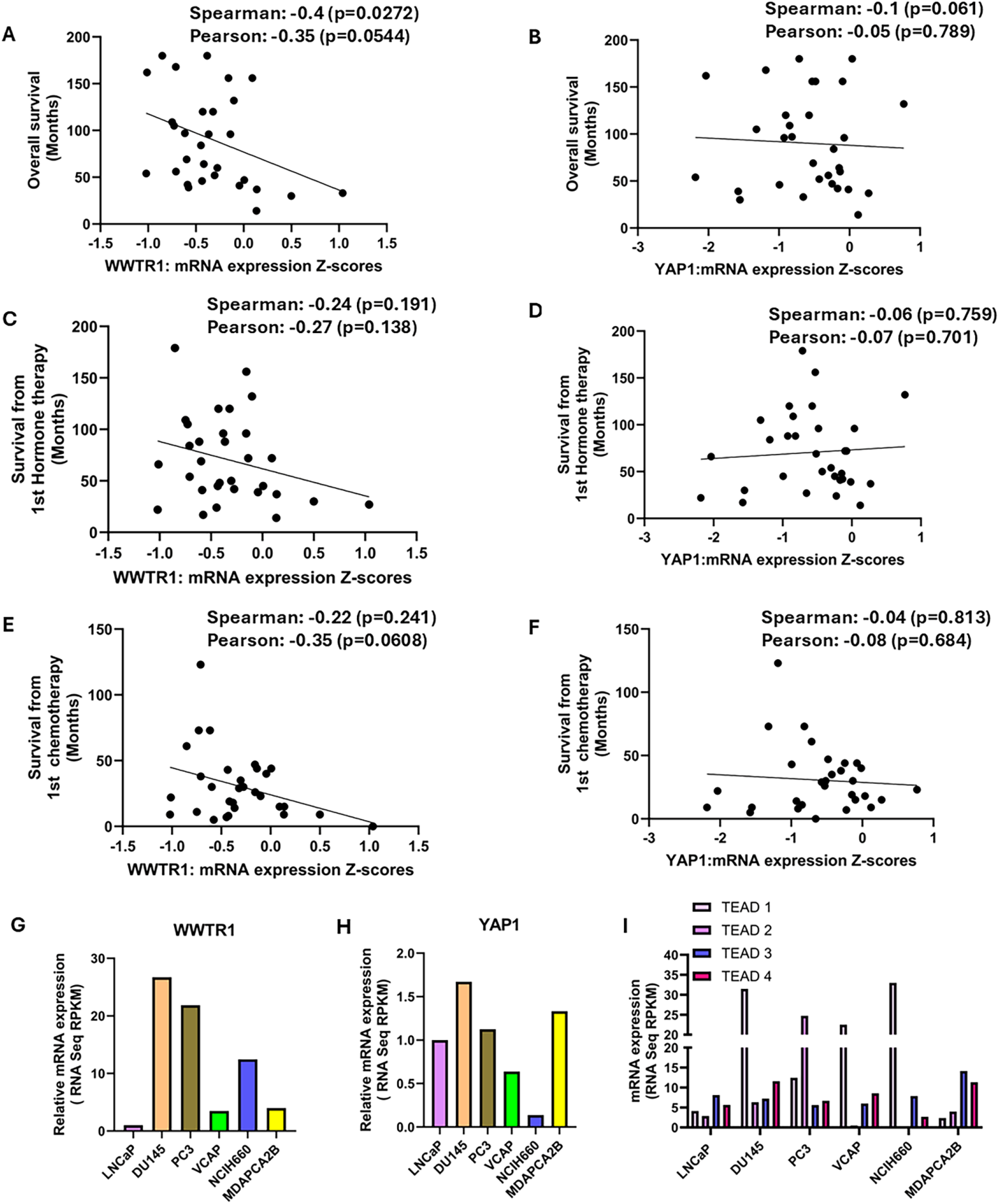
Elevated YAP1/WWTR1 associated with poor therapeutic response. Our analysis of clinical data (Metastatic Prostate Adenocarcinoma (MCTP, Nature 2012), 112 samples/119 patients) shows an inverse correlation between WWTR1 and YAP1 expression and overall survival (A, B), survival from 1st hormone therapy for WWTR1 and YAP1 (C, D), and survival from 1st chemotherapy (E, F), respectively. Moreover, YAP1 (G), WWTR1 (H), and TEAD (I) are differentially expressed across PCa cell lines. Data were downloaded from *cBioportal* (Next Generation Characterization of the Cell Line Encyclopedia; Ghandi et al., 2019).

## Discussion

Therapeutic resistance remains a major clinical challenge in advanced prostate cancer (PCa), especially after prolonged treatment with second-generation androgen receptor (AR)-directed inhibitors ^36^ . Although enzalutamide initially provides substantial clinical benefit, many patients eventually develop resistance, highlighting the urgent need to identify alternative, AR-independent survival pathways ^9^ . In this study, we examined the therapeutic potential of targeting the Hippo-YAP-1/TAZ-TEAD pathway across diverse advanced PCa models, focusing on enzalutamide-resistant disease. Our findings show that the small molecule CA3 potently suppresses key malignant phenotypes by pharmacologically disrupting the YAP-1/TAZ-TEAD transcriptional interface, including real-time proliferation, long-term clonogenic survival, migration, and transwell invasion. Importantly, these anti-tumor effects persist in independently derived, isogenic enzalutamide-resistant (ENZR) LNCaP and C4-2 cell models.

A major finding of this study is CA3’s potency in enzalutamide-resistant populations. To determine whether this therapeutic vulnerability shows a biological class effect of Hippo pathway inhibition rather than a compound-specific artifact, we also tested two structurally diverse small-molecule YAP-1/TAZ-TEAD inhibitors, GNE-7883 and K-975, in cell growth assays. Both compounds produced similarly potent anti-proliferative effects across our models (data not shown), strongly supporting the conclusion that advanced and treatment-resistant PCa cell states depend on functional YAP-1/TAZ-TEAD transcription.

While LNCaP ENZR and C4-2 ENZR cells continued to proliferate robustly and form colonies under continuous enzalutamide pressure, pharmacological pathway inhibition overcame this resistance. CA3 reduced not only baseline growth but also traits closely associated with metastatic potential, including migration and transwell invasion. These findings correspond with growing evidence that aberrant YAP-1/TAZ activation contributes to adaptive resistance to targeted cancer therapies ^37^ . Pathological upregulation of this pathway can activate alternative transcriptional networks and promote lineage plasticity, enabling tumor cells to survive despite maximal inhibition of their primary oncogenic drivers ^38,39^ . In advanced PCa, an AR-V7/YAP-11/TAZ signaling loop has been identified as a key driver of enzalutamide resistance ^40^ , supporting a functional, compensatory interaction between AR signaling variants and Hippo pathway effectors ^41^ .

Importantly, our data suggest that enzalutamide-resistant cells acquire a targetable dependence on downstream YAP-1/TAZ-TEAD signaling, rather than regaining baseline sensitivity to enzalutamide. This distinction is critical: resistant tumors frequently evade therapy by bypassing the AR axis through alternative transcriptional networks, rather than simply becoming insensitive to the drug ^17^. Disrupting these compensatory escape routes is therefore a rational strategy for eliminating persistent tumor populations that emerge during frontline AR-directed therapy ^41^ . Similar adaptive mechanisms have been reported in other solid tumors, where YAP-1/TAZ-TEAD activation enables escape from targeted kinase inhibitors and pharmacological TEAD disruption restores tumor control ^42^ . Thus, the YAP-1/TAZ-TEAD axis appears to act as a general survival mechanism that becomes increasingly dominant as tumors evolve under therapeutic selective pressure ^39,43^ .

Crucially, CA3’s anti-tumor efficacy extended beyond enzalutamide-resistant models. The compound significantly inhibited proliferation, clonogenicity, migration, and invasion in LNCaP, C4-2, DU145, and PC3 cell lines. A limited study using LNCaP and PC3 cells recently correlated these findings ^44^ . These broad effects suggest that YAP-1/TAZ-TEAD signaling regulates several classic hallmarks of aggressive PCa, including sustained proliferation, long-term survival, and cellular motility. Minor differences in drug sensitivity among the cell lines may reflect baseline pathway activity, AR status, or distinct genomic backgrounds. Nevertheless, CA3’s ability to suppress both AR-positive and AR-independent models shows its potential use across multiple stages and molecular subtypes of advanced disease.

Our novel in vivo findings further support the translational potential of these observations. Systemic CA3 administration significantly reduced final tumor weights in both DU145 and PC3 subcutaneous xenograft models. This anti-tumor response occurred without animal weight loss or adverse behaviors, indicating that the alternate-day dosing regimen (1.5 mg/kg) was well tolerated. Histopathological analysis showed clear structural changes in CA3-treated tumors, including increased collagen deposition, as shown by Masson’s Trichrome staining. Although the precise biological effects of this stromal remodeling call for further study, these changes suggest that inhibiting YAP-1/TAZ-TEAD signaling affects not only tumor parenchymal cells but also tumor-stroma interactions as well as extracellular matrix (ECM) assembly ^45,46^ . These observations are consistent with prior reports of CA3’s preclinical potency in other aggressive malignancies, including esophageal adenocarcinoma ^47^ and malignant mesothelioma ^48^ .

Mechanistically, immunohistochemical profiling confirmed that the documented reduction in tumor burden was driven by a concurrent decline in cell proliferation and an increase in programmed cell death. CA3-treated xenografts had significantly lower PCNA and Ki67 proliferative indices, together with a marked increase in fluorescent TUNEL-positive apoptotic cells. These in vivo findings are consistent with the established role of the YAP-1/TAZ-TEAD complex in regulating pro-survival and proliferative gene programs and confirm that disrupting this pathway compromises overall tumor maintenance ^25,49^ .

An intriguing finding was the striking accumulation of nuclear γ-H2AX foci following CA3 exposure, indicating activation of DNA damage signaling cascades. TEAD family proteins have been linked to DNA damage response regulation through direct physical interactions with core double-strand break repair proteins, including XRCC5, XRCC6, PARP1, and RIF1 ^50^. Accordingly, depleting or disrupting TEAD factors may impair homologous recombination and non-homologous end-joining pathways, resulting in persistent, unrepaired DNA lesions ^51,52^ . Although these insights yield a plausible account of the observed γ-H2AX accumulation, γ-H2AX can also accumulate downstream of general replication stress or during the active execution of apoptosis ^53^ . Future studies using complementary approaches, such as neutral comet assays or immediate DNA repair reporter assays, will be essential to determine whether CA3 directly disables the double-strand break repair machinery.

The convergence of DNA damage accumulation and apoptotic cascade activation highlights promising opportunities for rational combination therapies. If YAP-1/TAZ-TEAD inhibition fundamentally compromises a tumor cell’s ability to resolve genotoxic stress, pairing TEAD-directed agents with therapies that introduce DNA lesions or exploit repair defects could produce strong therapeutic synergy. In advanced PCa, this approach could be highly relevant when combined with poly(ADP-ribose) polymerase (PARP) inhibitors, though formal combination-scheduling studies are needed to validate this strategy ^53,54^ . Similarly, because CA3, GNE-7883, and K-975 effectively target enzalutamide-resistant cells, administering YAP-1/TAZ-TEAD inhibitors alongside or immediately after frontline AR-directed blockades may inhibit the emergence of resistant subclones. Future combination studies featuring formal synergy modeling (e.g., Bliss independence or Chou-Talalay methods) will be necessary to distinguish true restoration of therapeutic sensitivity from simple additive effects.

The clinical and biological relevance of this axis is supported by expression analyses in a prostate cancer patient cohort and representative cell lines. A previous study has indicated that YAP1 overexpression contributes to the development of enzalutamide resistance by induction of cancer stemness and lipid metabolism in prostate cancer ^55^ . Our analysis found that elevated tumor expression of WWTR1 (encoding TAZ) and, to a lesser extent, YAP-1 inversely correlates with overall survival, with a trend toward shorter survival after frontline hormone and chemotherapy treatments. In parallel, our cell line panel showed heterogeneous expression of WWTR1, YAP-1, and TEAD1–4 across PCa models. Notably, baseline WWTR1 transcript levels were markedly elevated in aggressive, AR-independent cell lines such as DU145 and PC3 and the neuroendocrine cell line NCIH660. This variation indicates that cellular dependence on active YAP-1/TAZ-TEAD signaling depends on the tumor’s specific molecular condition ^56^ rather than being uniform across all prostate malignancies. Taken together, these data show that YAP-1/TAZ-TEAD are clinically relevant targets and suggest that WWTR1 is a potential prognostic indicator of therapy resistance in aggressive prostate cancer.

In conclusion, our study offers strong preclinical evidence that the YAP-1/TAZ-TEAD signaling axis represents a targetable therapeutic vulnerability in advanced prostate cancer, including tumors that have escaped enzalutamide control. Pharmacological disruption of this axis with CA3 consistently suppressed aggressive malignant phenotypes in vitro and significantly restricted tumor growth in vivo in both AR-dependent and AR-independent models. Along with the novel finding that treatment induces DNA damage accumulation, these data strongly support the development of YAP-1/TAZ-TEAD-targeted strategies to counter therapy resistance and lineage plasticity in patients with advanced prostate cancer.

## Author Contributions (CRediT Taxonomy)

• **Vandana Mohan:** Methodology, Validation, Formal analysis, Investigation, Data curation, Writing - original draft, Visualization.

• **Santosh Lamichhane:** Investigation, Data acquisition, Analysis.

• **Praveen Kumar Jaiswal:** Investigation, Data Analysis, Writing-review C editing.

• **Sweaty Koul:** Conceptualization, Methodology, Investigation, Management, Writing -review C editing.

• **Guobin Kang:** Image acquisition, methodology, and investigation.

• **Dorota D. Wyczechowska:** Investigation, Resources (Flow Cytometry), Software.

• **Hari K Koul:** Conceptualization, Resources, Writing - review C editing, Supervision, Project administration, Funding acquisition.

---

**Figure S1:**
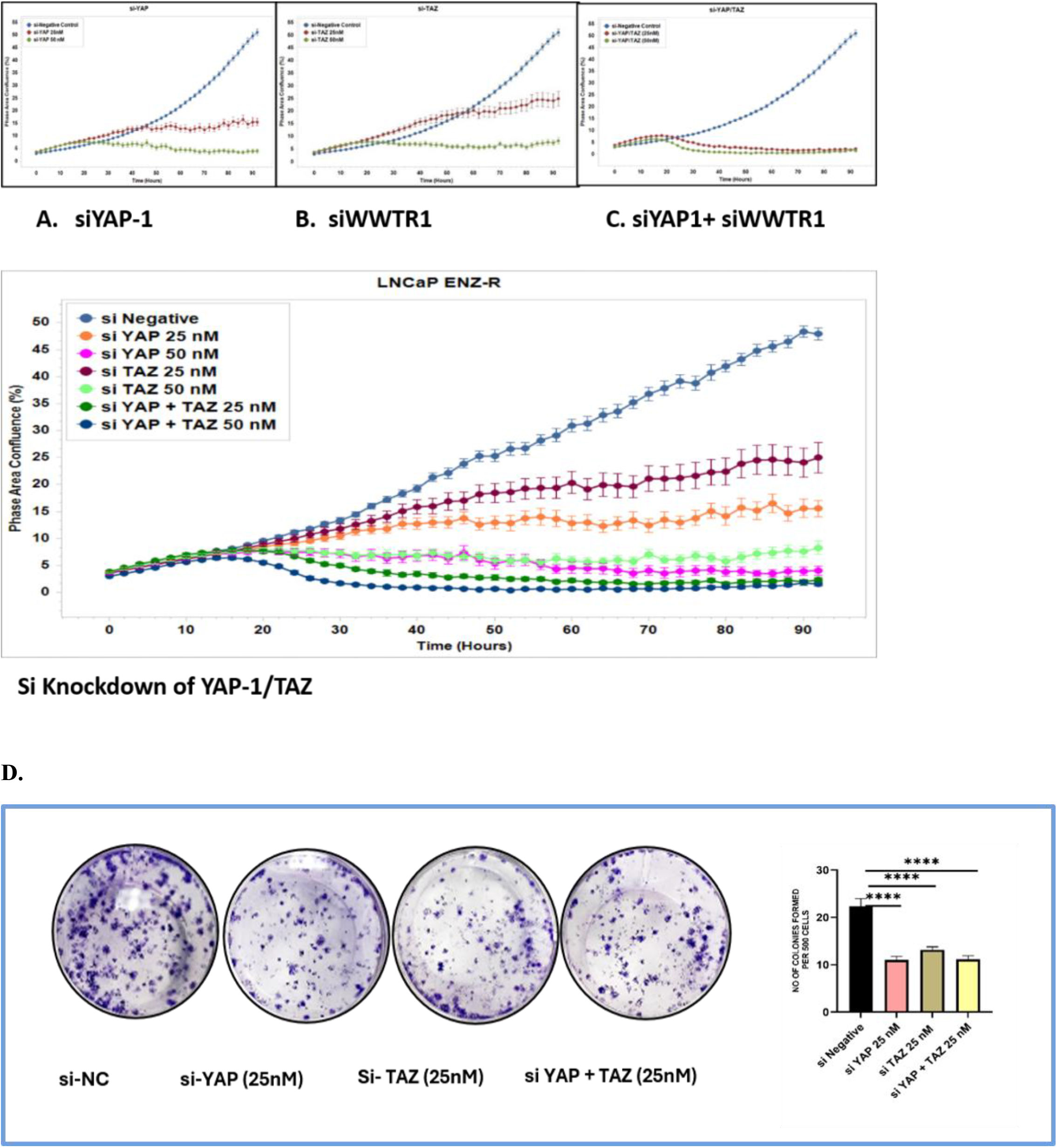
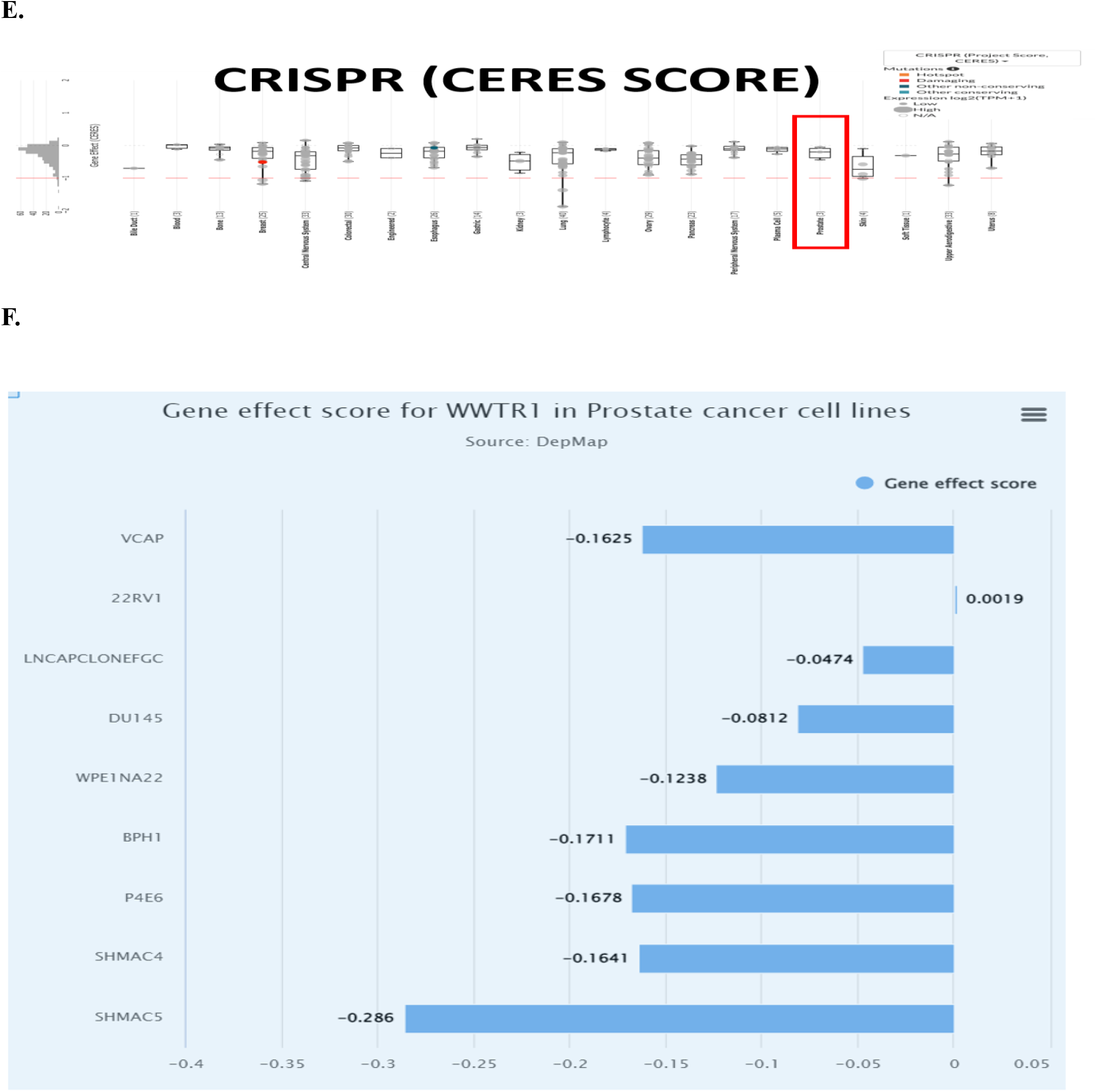
Transient depletion of YAP and TAZ using siRNA reduces cell proliferation and colony-forming ability in LNCaP enzalutamide-resistant prostate cancer (PCa) cells. Cell proliferation curves after si-YAP treatment at 25 and 50nM (A); Cell proliferation curves following si-TAZ treatment at 25 and 50nM (B); Combined si-YAP/TAZ treatment at 25 and 50nM in LNCaP ENZ-R cells (C). These data are split from the figure below for clarity. Data are presented as mean ± SD (n=6). Colony formation assay after si-YAP/TAZ treatment at 25nM (D); Right panel: quantification of colony numbers per treatment group; Data are presented as Mean ± SD (n=3; **** p≤0.0001 compared with the respective controls). The CERES dependency score maps for WWTR1 (E,F) were plotted using the online DepMap portal (Broad Institute), showing that WWTR1 is indispensable for most of the prostate cancer cell lines

**Table S1:**
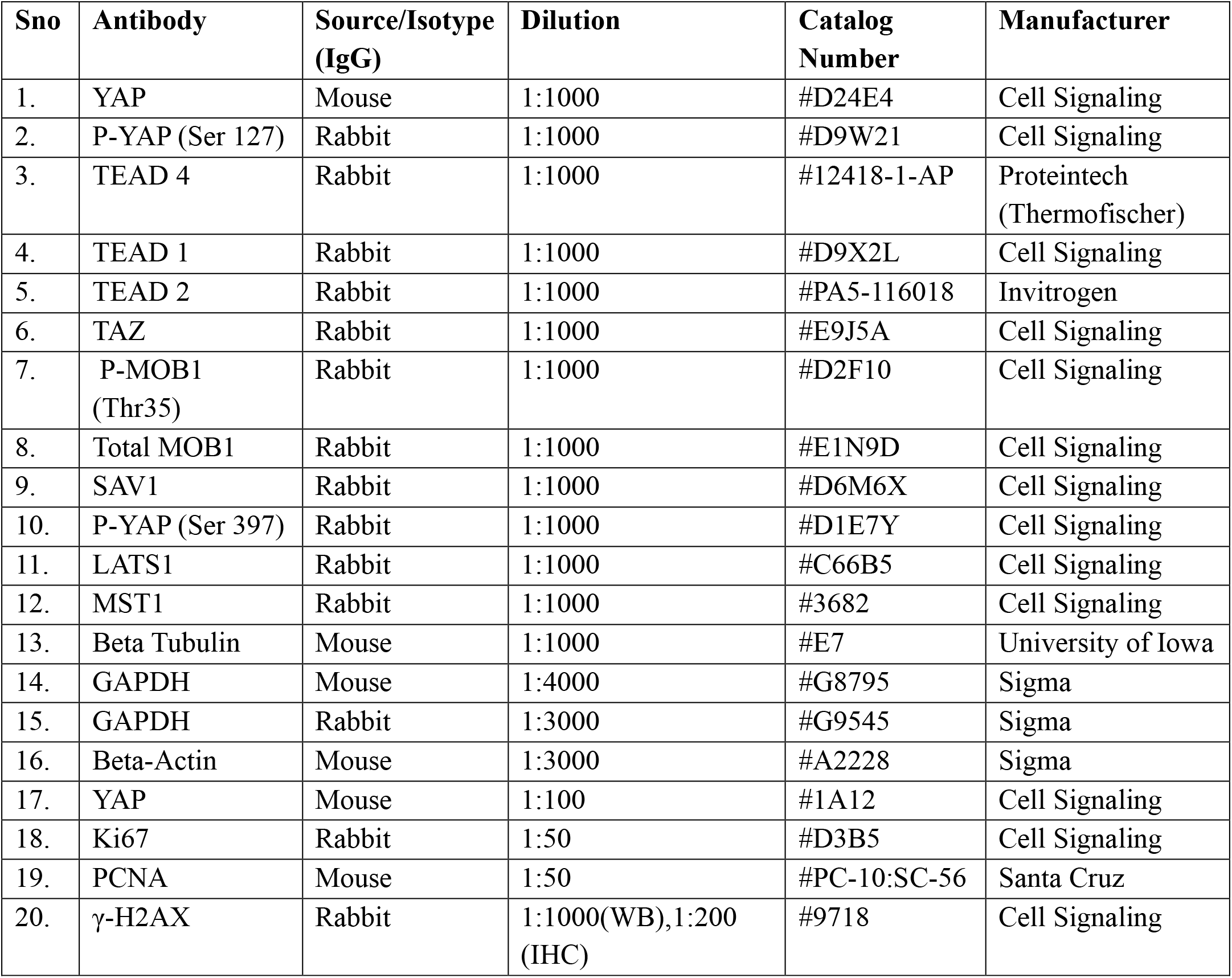
List of Primary Antibodies.

**Table S2:** List of Secondary Antibodies.

| **Sno** | **Antibody** | **Dilution** | **Catalog Number** | **Manufacturer** |
|---|---|---|---|---|
| 1. | Anti-Rabbit IgG, HRP-linked antibody | 1:1000 | #7074 | Cell Signaling |
| **2.** | Anti-Rabbit IgG, HRP-linked antibody | 1:5000 | #111-035-003 | Jackson Laboratory |
| **3.** | Anti-Mouse IgG, HRP-linked antibody | 1:5000 | #115-035-003 | Jackson Laboratory |

**Table S3:**
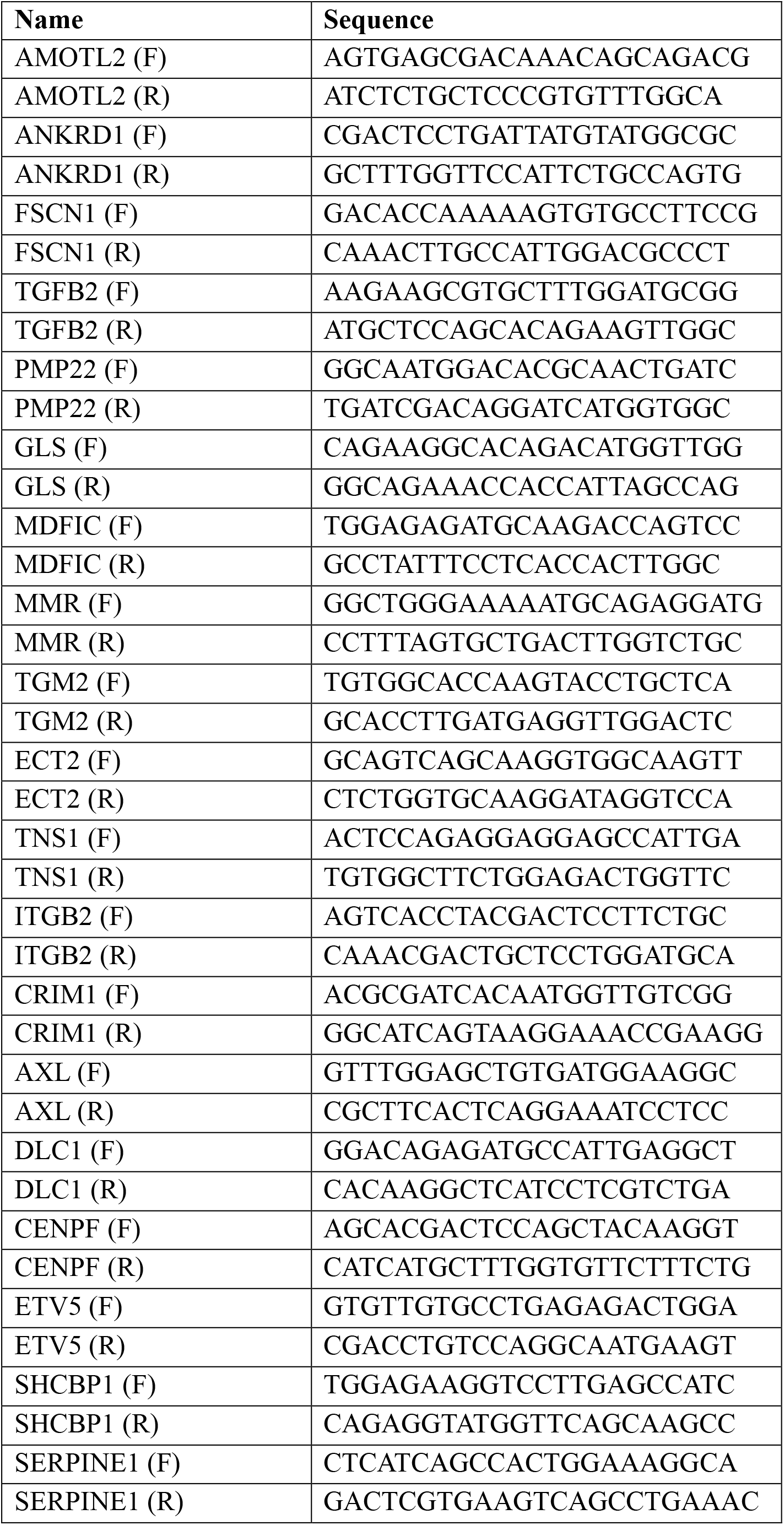
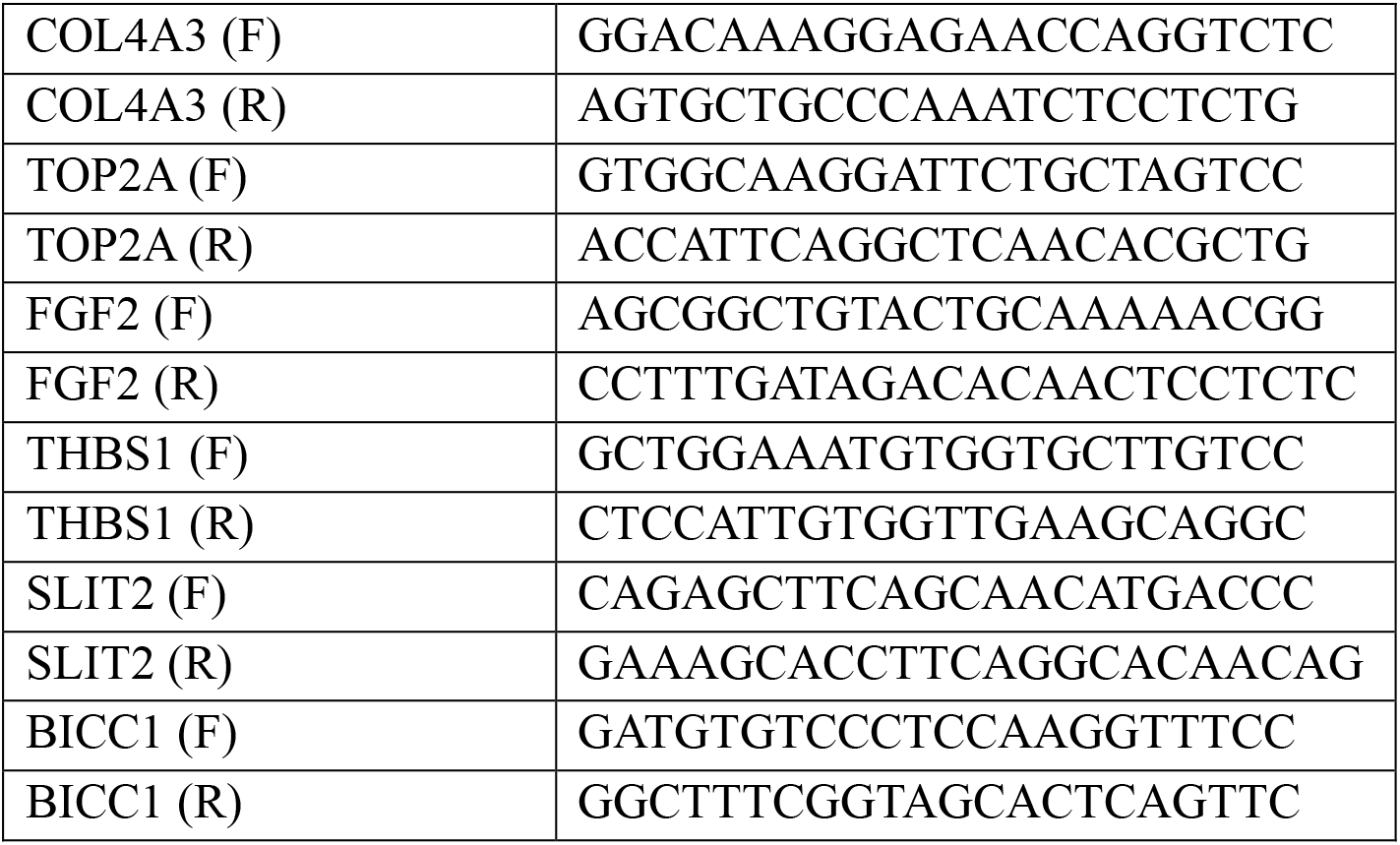
List of Primers.

## Supplementary document

**Figure1 B.**
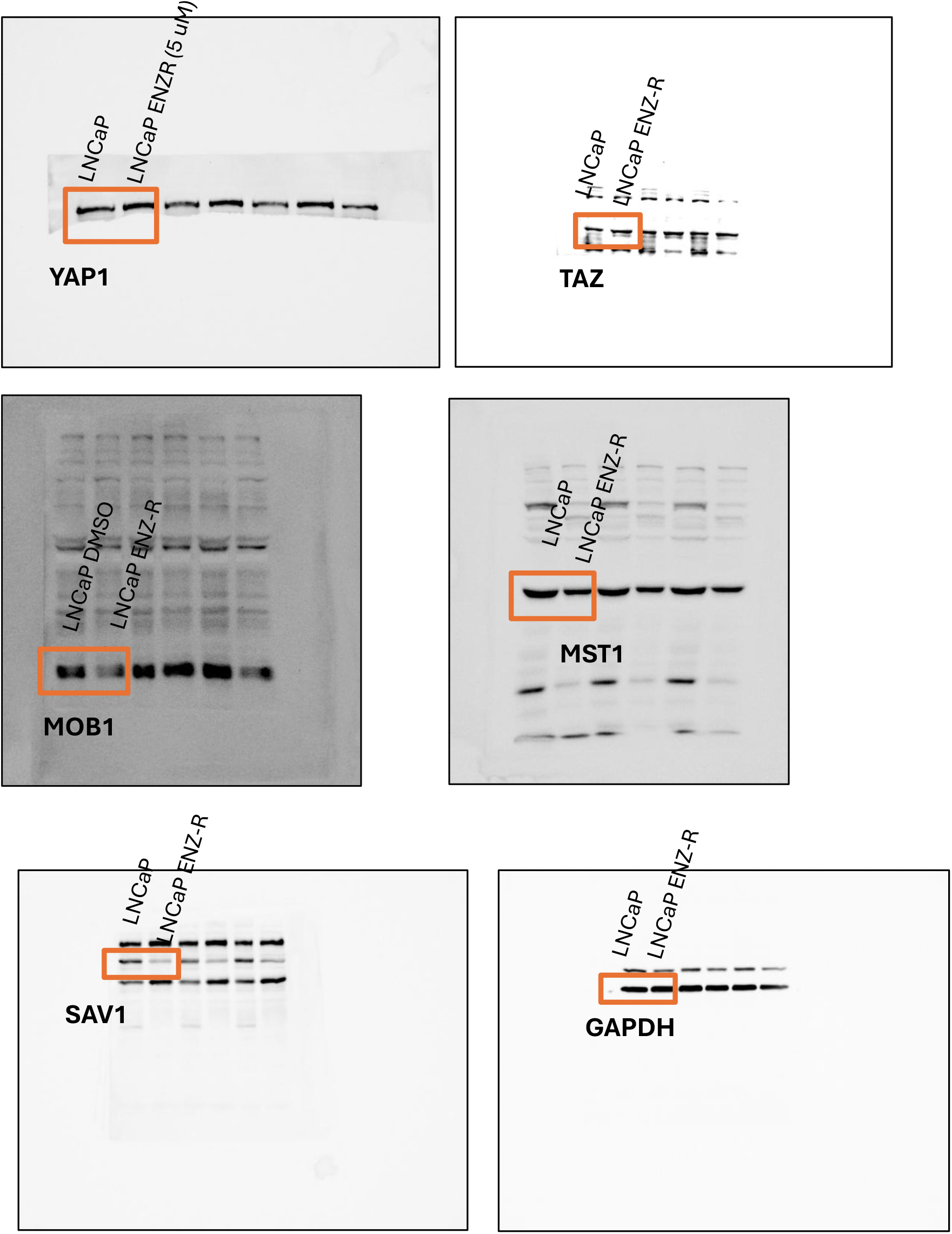
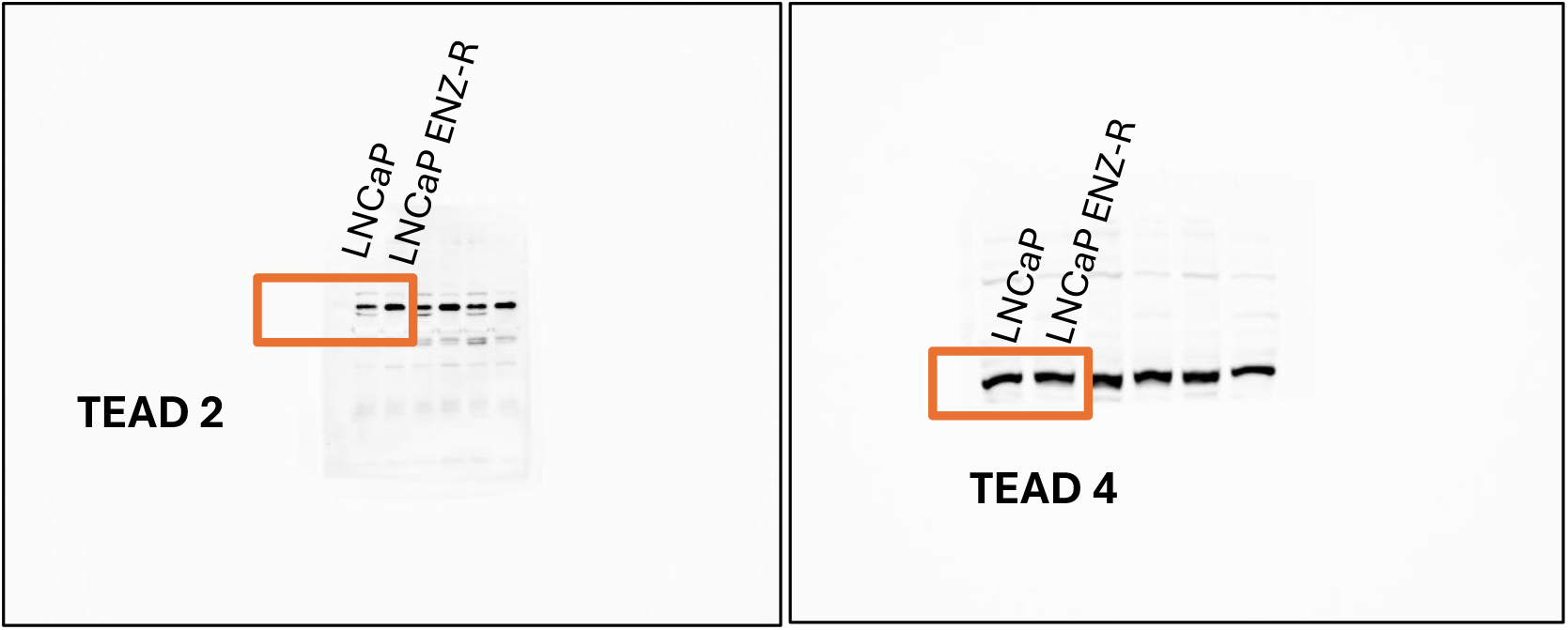

**Figure 1 C.**
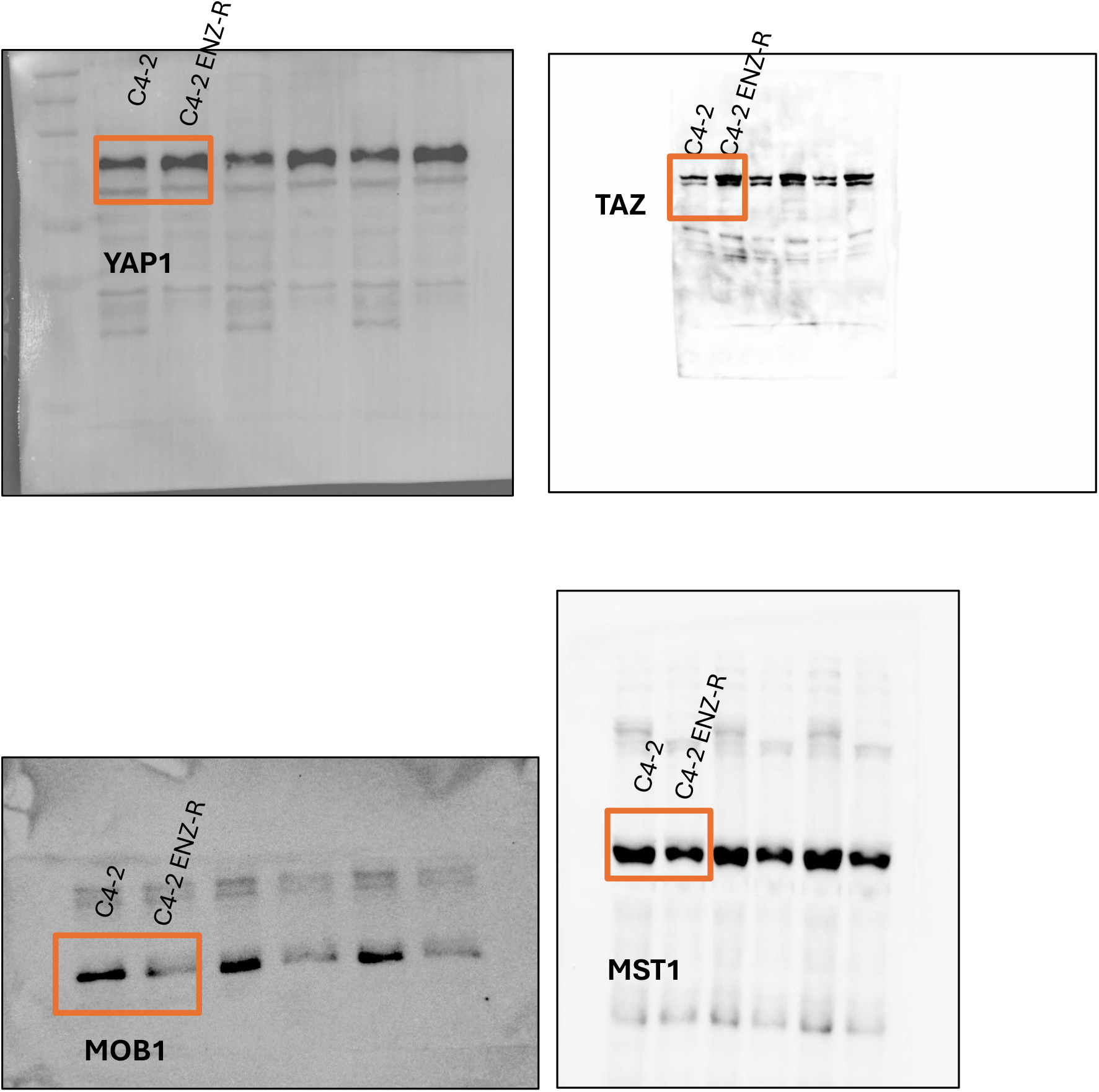
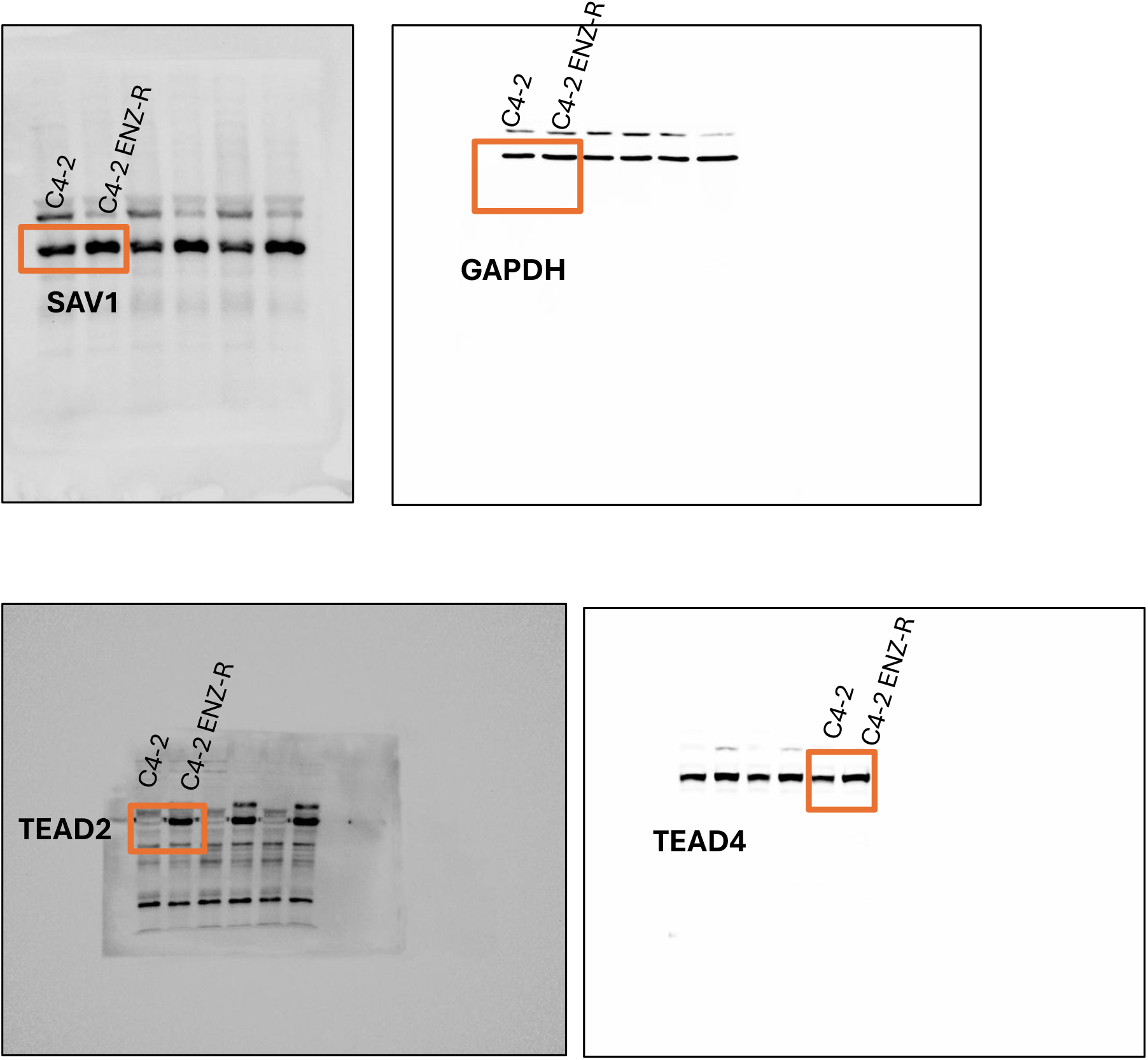

**Figure8 E.**
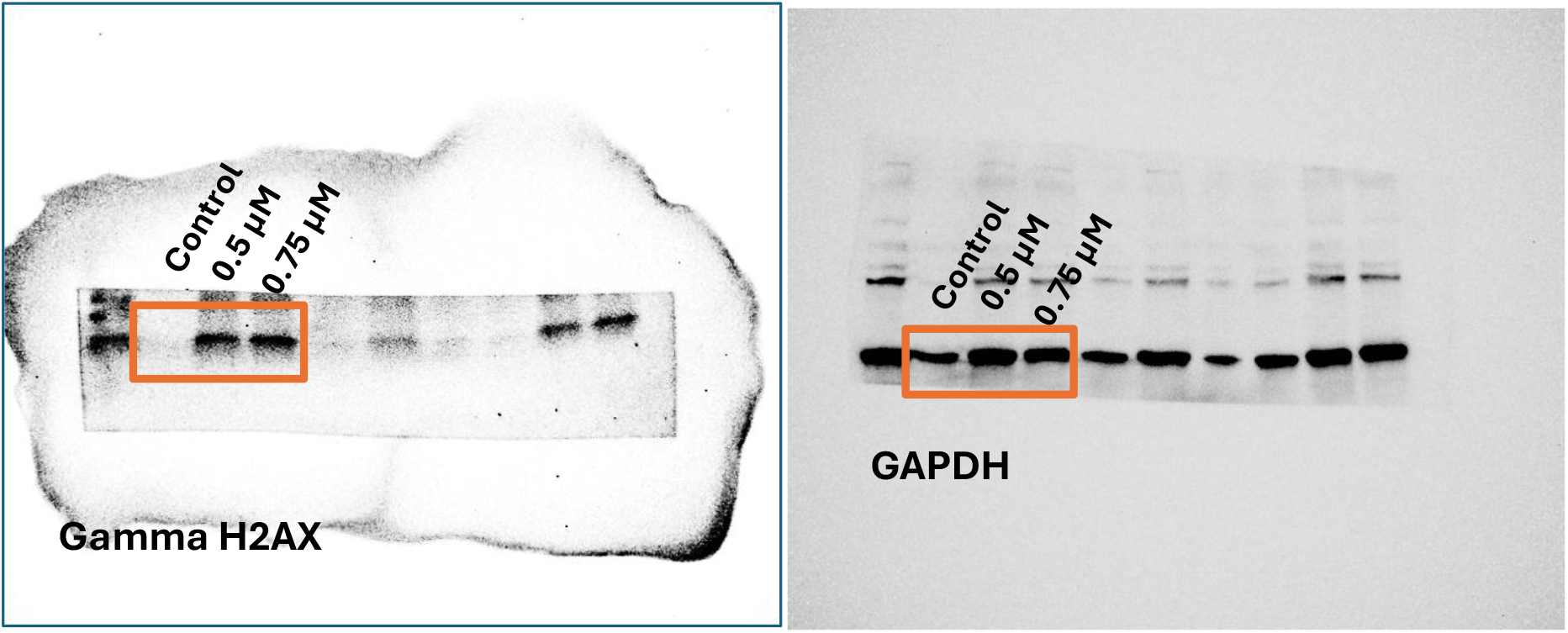
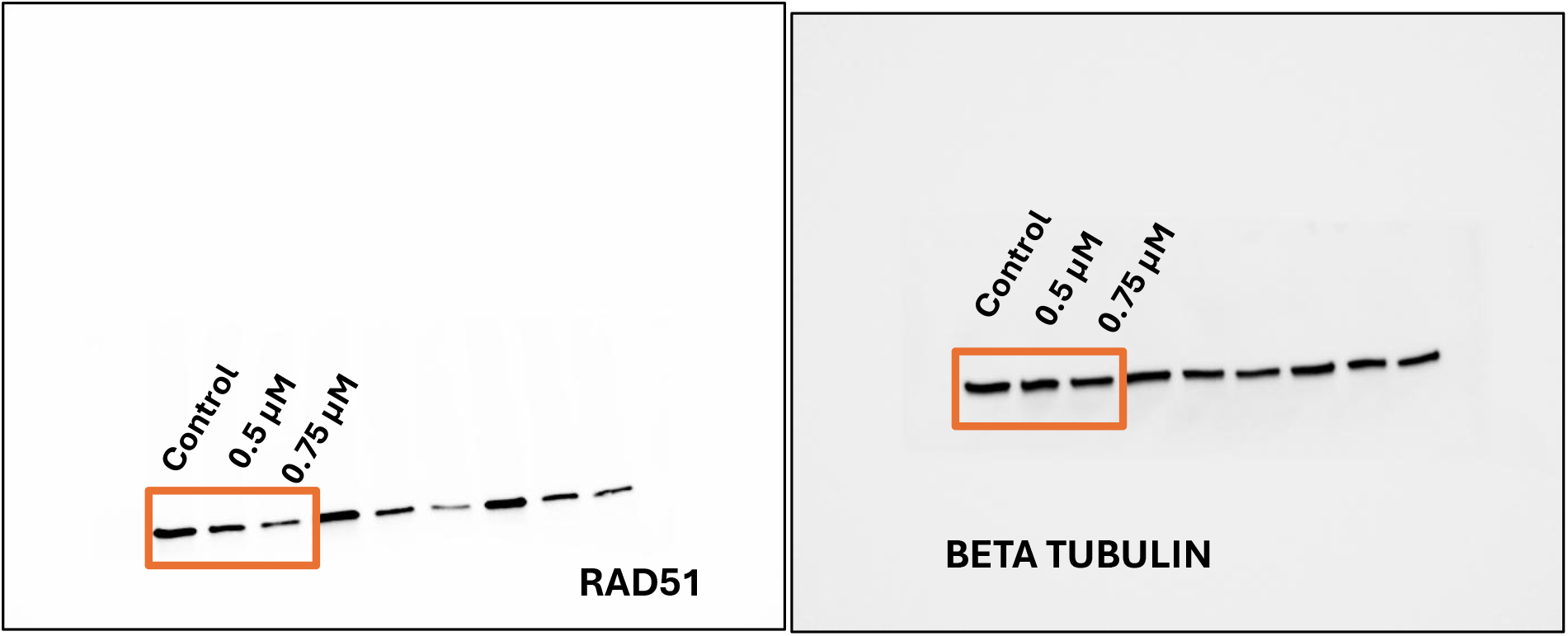
DU145

**Figure. 8H.**
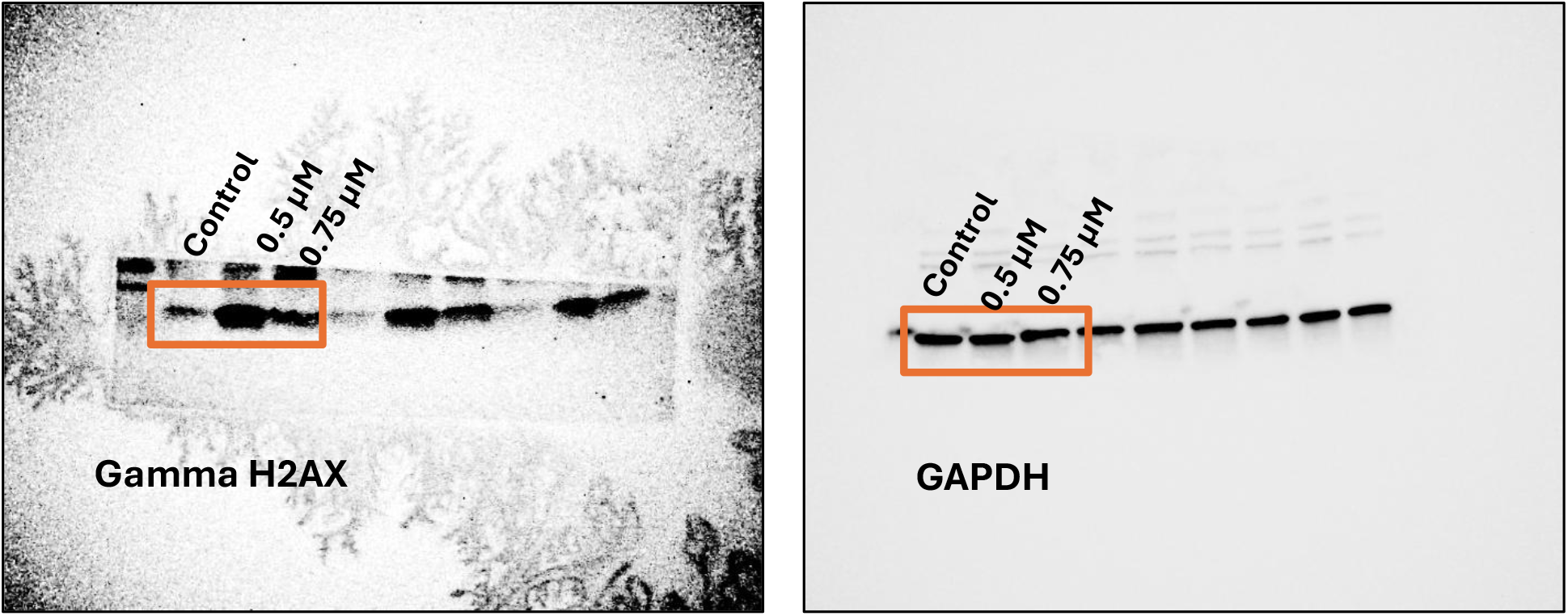
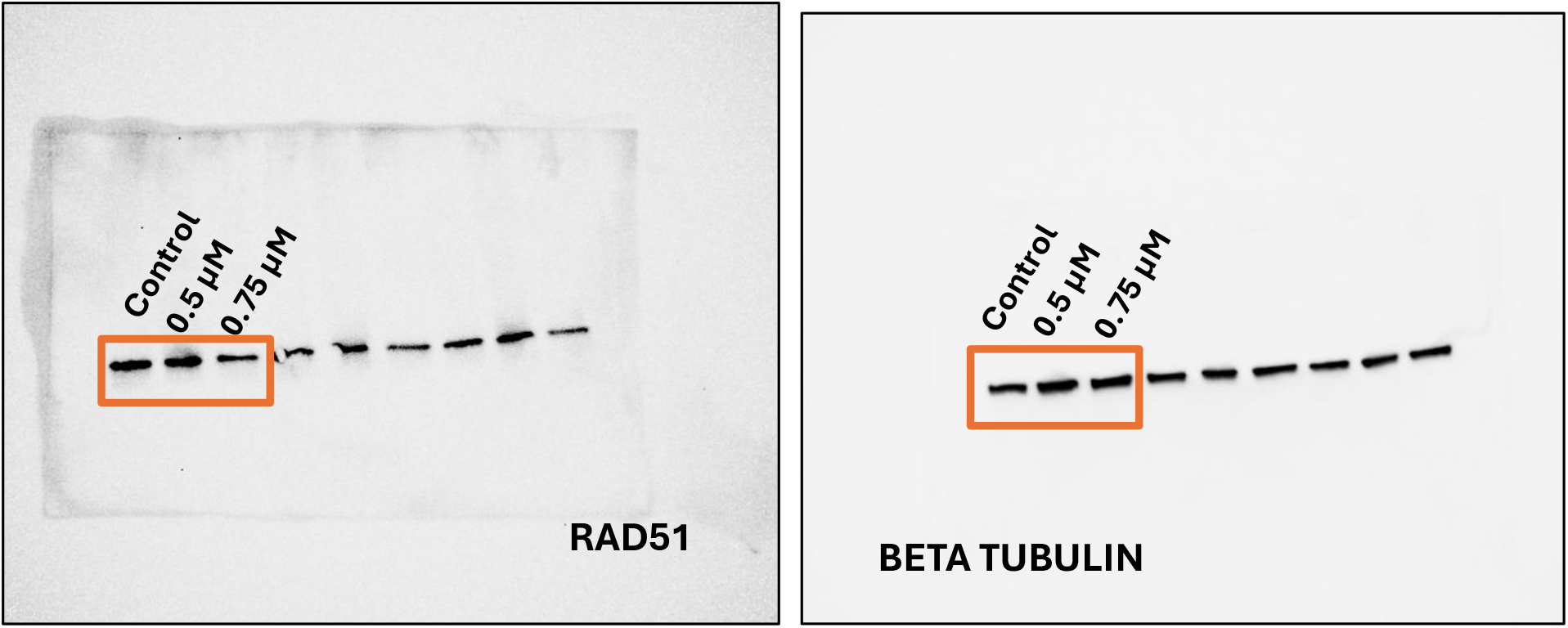
PC3

